# Deaminase-mediated chromatin accessibility profiling with single-allele resolution

**DOI:** 10.1101/2024.12.17.628768

**Authors:** Tian Yu, Zhijian Li, Ellie Gibbs, Reina Iwase, Matthew J Francoeur, Quang vinh Phan, Jing Zhao, Jane Rosin, Phillip A Cole, Luca Pinello, Richard I Sherwood

**Author notes:** Corresponding author. (L.P.), (R.I.S.). These authors contributed equally to this work (Tian Yu, Zhijian Li, Ellie Gibbs).

## Abstract

Binding of transcription factors (TFs) at gene regulatory elements controls cellular epigenetic state and gene expression. Current genome-wide chromatin profiling approaches have inherently limited resolution, complicating assessment of TF occupancy and co-occupancy, especially at individual alleles. In this work, we introduce Accessible Chromatin by Cytosine Editing Site Sequencing with ATAC-seq (ACCESS-ATAC), which harnesses a double-stranded DNA cytosine deaminase (Ddd) enzyme to stencil TF binding locations within accessible chromatin regions. We optimize bulk and single-cell ACCESS-ATAC protocols and develop computational methods to show that the increased resolution compared with ATAC-seq improves the accuracy of TF binding site prediction. We use ACCESS-ATAC to perform genome-wide allelic occupancy and co-occupancy imputation for 64 TFs each in HepG2 and K562, revealing that the propensity of a majority of TFs to co-occupy nearby motifs oscillates with a period approximating the helical turn of DNA. Altogether, ACCESS-ATAC expands the resolution and capabilities of bulk and single-cell epigenomic profiling.

## Main text

Human tissue specialization and disease states are controlled by the epigenetic state of cells, which is largely determined by the actions of over 1,600 sequence-specific transcription factors (TFs)^1^. TFs recruit the machinery that controls gene expression and long-term gene regulatory memory. The large number of TFs, combined with their complex hierarchical and combinatorial binding logic^2^, has hindered a predictive understanding of how TFs bind to target sites across cell states.

Measurement of the accessibility of chromatin to DNA-modifying enzymes^3^ has proven valuable in mapping gene regulatory elements and identifying their component TF binding sites^4,5^. However, current methods of genome-wide chromatin accessibility assessment have deficiencies that motivate new approaches.

In the most common chromatin accessibility measurement techniques, DNase-seq^6^ and ATAC-seq^7^, enzymatic cleavage or Tn5 transposition, respectively, is used to map the relative accessibility of nucleotides across the genome. The majority of gene regulatory elements show heightened accessibility^4^. Within such regions, TF binding creates local occlusions, or footprints, which can be used to infer TF binding sites across the genome^4,8,9^.

These approaches have two inherent limitations. First, because accessibility measurement depends on termination of a DNA strand, single or paired end sequencing reads can provide at most two data points, from the two ends of the fragment. Second, these ends must be separated by at least 40 nucleotides in order to reliably map the resulting fragment to a reference genome. These limitations inherently constrain the resolution of DNase-seq and ATAC-seq. They prevent analysis of co-occurrence of adjacent TF binding or nucleosome positioning events at an allelic level. Additionally, and crucially, while ATAC-seq can be scaled down to single-cell level (scATAC-seq)^10^, its resolution is inherently bounded by sparsity, precluding assessment of TF binding and nucleosome positioning in single cells. Instead, pseudo-bulk analysis, in which dozens to hundreds of similar cells are combined, is used to infer regulatory state of populations from scATAC-seq data^11^.

Several distinct chromatin accessibility measurement approaches have been developed that do not rely on strand termination and thus circumvent the limitations of DNase-seq and ATAC-seq. In Fiber-seq and SMAC-seq^12,13^, chromatin is treated with a DNA N^6^-adenine methyltransferase, and long-read sequencing is used to distinguish methylated and unmethylated adenines. These methods allow for single-molecule stenciling of chromatin accessibility, revealing TF binding footprints at the allelic level. However, they rely on long-read sequencing platforms and are incompatible with PCR amplification due to erasure of the methylation, rendering them expensive for high-resolution genome-wide data and currently incompatible with single-cell application. In NOMe-seq/single-molecule footprinting (SMF)^14,15^, chromatin is treated with a GpC methyltransferase, and either long-read sequencing^16^ or bisulfite conversion is used to provide single-molecule chromatin accessibility profiling. When used with long-read sequencing, NOMe-seq/SMF has the same drawbacks as Fiber-seq. When paired with bisulfite, the use of a GpC methylase limits resolution to ∼1/8 of nucleotides, bisulfite-induced DNA degradation limits sensitivity, and the lack of a simple approach to enrich reads in accessible chromatin makes it challenging to achieve high coverage in active chromatin. Altogether, there is currently no genome-wide approach ideally suited to high-resolution, single-cell and single-molecule chromatin accessibility profiling.

Double-stranded DNA cytosine deaminase (Ddd) enzymes are a newly discovered class of enzymes that catalyze cytosine to uracil editing. The first discovered double-stranded DNA cytosine deaminase, DddA, natively acts as an inter-bacterial toxin and has been engineered into a tool for mitochondrial genome editing^17^. DddA has strong preference for editing at TC motifs, but enzyme engineering and ortholog mining efforts have identified Ddd enzymes with relaxed sequence preferences ^18–21^. Aside from uses in mitochondrial editing, DddA has been fused with TFs in bacteria to map protein-DNA interaction sites^22^.

We reasoned that Ddd editing might improve the resolution of chromatin accessibility profiling due to selective editing activity. Cytosine deamination events can be detected through PCR-based conversion of uracil to thymine, making Ddd treatment compatible with all sequencing workflows. In this work, we show that Ddd enzymes exhibit chromatin accessibility-dependent cytosine deamination genome-wide. We develop an approach, Accessible Chromatin by Cytosine Editing Site Sequencing with ATAC-seq (ACCESS-ATAC), in which we treat chromatin concurrently with Ddd enzyme and Tn5 transposase. ACCESS-ATAC uses the same workflow as ATAC-seq, allowing readout by short-read sequencing and retaining the enrichment of accessible chromatin provided by ATAC-seq while also providing allele-level footprinting through detection of Ddd deamination events. We find that ACCESS-ATAC improves TF binding site prediction as compared with ATAC-seq. Moreover, ACCESS-ATAC enables allele-level TF occupancy and co-occupancy imputation across the accessible genome. Through imputation of allelic co-occupancy patterns of 64 TFs in K562 and HepG2 at over 1.5 million adjacent motif pairs, we find that the vast majority of TF pairs co-occupy nearby binding sites more often than expected under a model of independent binding. Moreover, we find widespread ∼10-nt periodicity in the co-occupancy of many human TFs, revealing a previously underappreciated logic underlying TF binding site choice. We further develop a single-cell ACCESS-ATAC workflow that improves resolution of TF footprint detection in single cells, enabling the generation of equivalent quality TF footprints using 2-3-fold fewer cells. Altogether, ACCESS-ATAC provides a flexible platform for high-resolution chromatin accessibility and TF binding profiling.

## Results

### Ddd enzymes induce chromatin accessibility-dependent cytosine deamination

To assess chromatin accessibility dependence of Ddd editing, we produced purified DddA^17^ (**Supplementary** Fig. 1) and titrated its activity in intact human HCT116 colorectal carcinoma nuclei at the accessible *PTMA* promoter and an inaccessible locus on chromosome 4^23^. Loci were PCR-amplified using Q5U polymerase, which converts uracils to thymines, and editing efficiency was measured using nextgen sequencing (NGS). Editing was dose-dependent, and the *PTMA* promoter had substantially higher editing than the inaccessible locus, primarily at known DddA target TC motifs (**Supplementary** Fig. 2). We next performed whole genome sequencing (WGS) after DddA treatment of HCT116 nuclei, finding that TC to TT edit fraction steadily increased in concert with published HCT116 ATAC-seq chromatin accessibility (**Supplementary** Fig. 2). Altogether, these data indicate that DddA provides chromatin accessibility-dependent cytosine deamination. Going forward, we refer to the treatment of intact nuclei with Ddd enzyme as Accessible Chromatin by Cytosine Editing Site Sequencing (ACCESS).

DddA’s preferential editing at TC dinucleotides limits its chromatin accessibility measurement resolution. We evaluated six Ddd enzymes reported to have low motif specificity^19,21^ through *in vitro* transcription/translation of each enzyme followed by ACCESS-WGS of intact HCT116 nuclei. We observed that, for samples treated with less motif-specific Ddd enzymes, the alignment of reads to the genome using the standard Bowtie2^24^ parameters decreased as chromatin accessibility increased, presumably due to the failed alignment of highly edited reads (**Supplementary** Fig. 3). We developed an alignment approach with iteratively lower mismatch penalties (**Methods**), which recovered reads from accessible regions of the genome while not substantially increasing alignment in Ddd-untreated WGS samples (**Supplementary** Fig. 3). Using this custom alignment, all six enzymes showed chromatin accessibility-dependent editing at certain trinucleotides; however, two highly homologous enzymes, the deaminases from *Simiaoa Sunii*, which we refer to as DddSs (also called Ddd1^19^, LbsDa01^21^, and SsDddA^25^) and a highly homologous, metagenomically identified deaminase MGYPDa829^21^, showed the most consistently enriched editing across sequence contexts (**Fig. 1A**, **Supplementary** Fig. 2).

**Figure 1:**
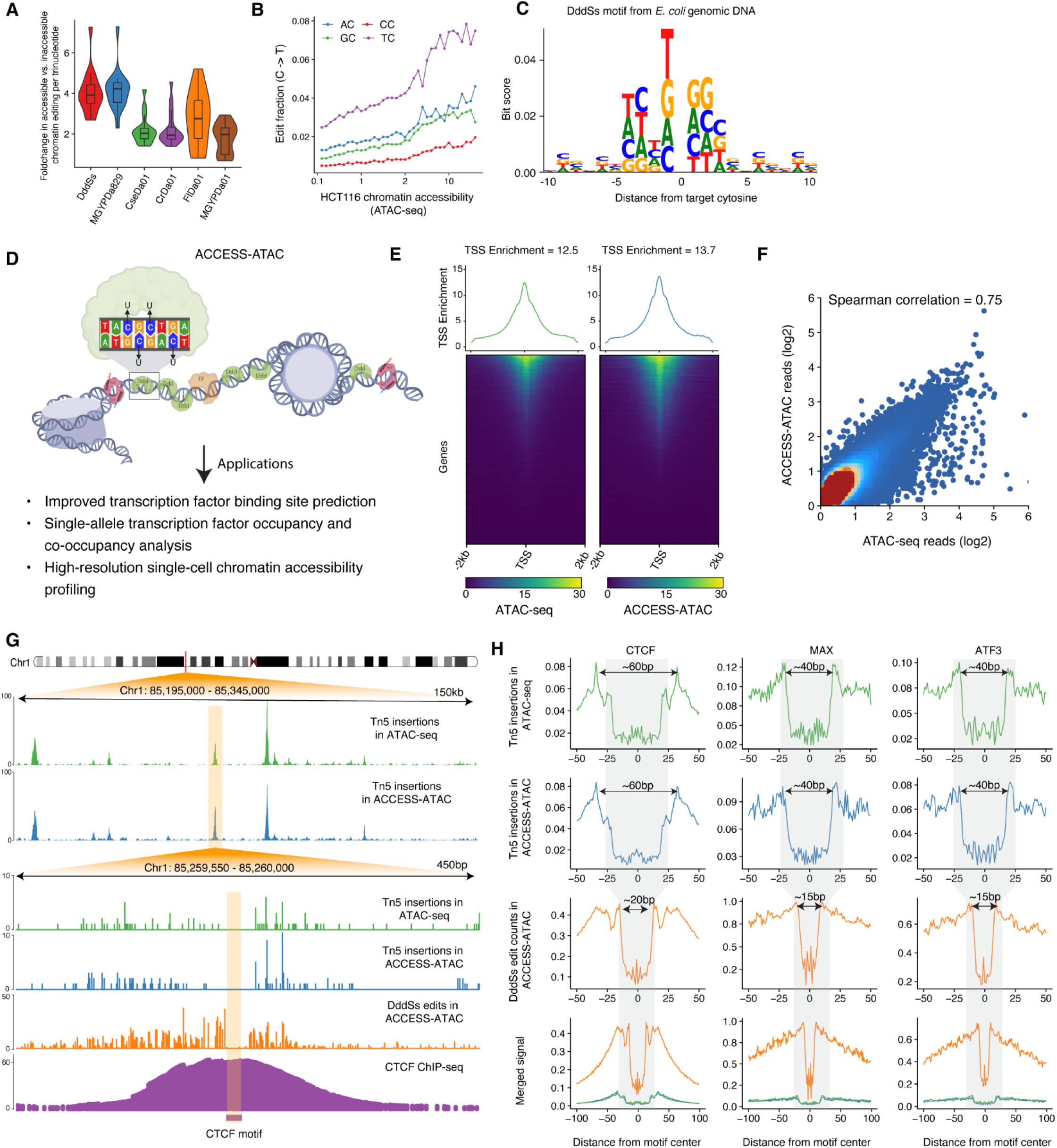
Chromatin accessibility-dependent cytosine deamination by Ddd proteins enables high-resolution transcription factor footprinting. (A) Enrichment of edited cytosines in accessible vs. inaccessible chromatin for 6 Ddd enzymes, separated by trinucleotide. Data is derived from ACCESS-WGS data, and each point represents the accessible vs. inaccessible foldchange in observed C2T editing for a particular cytosine-centered trinucleotide. (B) Dinucleotide-resolved edit fractions in DddSs-treated HCT116 cells binned by ATAC-seq coverage. Data derive from ACCESS-WGS data. (C) Motif logo showing the relative importance of each nucleotide surrounding the target cytosine in influencing DddSs editing efficiency in *E. coli* genomic DNA. (D) Schematic of ACCESS-ATAC, in which nuclei are concurrently treated with DddSs, which converts cytosine residues to uracils in proportion to chromatin accessibility, and Tn5. (E) Enrichment of transcription start sites (TSS) in K562 ATAC-seq and ACCESS-ATAC reads. Each row is a TSS, rows are ordered by the magnitude of read enrichment, and a summary plot above shows read enrichment in the 2 kb surrounding all TSS. (F) Scatter plot showing genome-wide correlation between ATAC-seq and ACCESS-ATAC coverage. Each dot represents a 10-kb bin, and the color refers to density of the bins. (G) Representative K562 ATAC-seq and ACCESS-ATAC signal tracks. Top: Tn5 insertion counts from K562 ATAC-seq (green) and ACCESS-ATAC (blue) in a representative 150-kb genomic region. Bottom: Nucleotide-level ATAC-seq Tn5 insertion counts (green), ACCESS-ATAC Tn5 insertion counts (blue) and DddSs edit counts (red), and smoothed K562 CTCF ChIP-seq counts and motif (purple) in a 450-bp subregion. Tn5 insertion and DddSs editing are depleted at the CTCF motif, and DddSs editing shows denser surrounding signal. (H) Metaplots showing K562 ATAC-seq Tn5 insertion signal (green), and ACCESS-ATAC Tn5 insertion (blue) and DddSs editing (orange) signal surrounding K562 ChIP-seq binding sites for CTCF, MAX and ATF3. Bottom plot shows merged non-normalized signal tracks to highlight the denser DddSs editing signal. Arrows denote footprint width.

We produced DddSs and its inhibitor (DddSs-I) through bacterial expression (**Supplementary** Fig. 1), performing ACCESS-WGS in HCT116 at an optimized dose of 250 nM. DddSs ACCESS-WGS data revealed a relatively permissive motif with considerably broader editing than DddA genome-wide (**Fig. 1B-C**). All further ACCESS experiments employed DddSs.

### High-resolution profiling accessible chromatin with ACCESS-ATAC

It is difficult to obtain the sequencing coverage needed for high-resolution dissection of TF binding using ACCESS-WGS since DNA fragmentation does not preferentially enrich for reads in accessible chromatin. We thus sought to combine ACCESS with ATAC-seq, using ATAC to focus sequencing depth on accessible chromatin and ACCESS to improve resolution within each sequenced allele. We compared two approaches to combine ACCESS with ATAC-seq (ACCESS-ATAC) in K562 cells, a cell line in which TF binding sites have been characterized extensively by ChIP-seq^17^. We tested 30-minute concurrent treatment of intact chromatin with Tn5 and DddSs (concurrent ACCESS-ATAC, **Fig. 1D**), as well as a three-step sequential protocol treating intact nuclei with DddSs for variable times (15-, 30-, and 60-minute), quenching DddSs activity with DddSs-I, and then performing 30-minute Tn5 treatment (sequential ACCESS-ATAC). We also compared the resulting data with 30- minute Tn5 treatment alone as an important baseline against the standard ATAC-seq protocol.

All approaches yielded NGS libraries with similar fragment diversity as measured by qPCR (**Supplementary** Fig. 4), suggesting that DddSs treatment does not substantially alter Tn5 integration or library amplification when using Q5U polymerase. We developed a metric of ACCESS data quality based on DddSs editing fraction surrounding CTCF ChIP-seq binding sites since CTCF binding profoundly alters chromatin accessibility and binds to a subset of common loci across cell types^26^. As expected, we found that ACCESS-ATAC data exhibited high editing in peak regions surrounding CTCF binding sites, low editing at the footprint where CTCF binding occludes editing, and a wave-like histone coordination pattern radiating distally from binding sites (**Supplementary** Fig. 4). We thus determined the signal strength of ACCESS data by measuring the DddSs edit fraction (C to T and G to A) in the peak regions and the signal-to-noise as the log-fold change in editing between the peak and valley regions. We observed that 30-minute concurrent and sequential ACCESS-ATAC yielded equivalent ACCESS signal strength (CTCF peak edit fraction) and signal-to-noise (peak/valley LFC), while shortening the DddSs treatment time below 15 minutes decreased signal strength and signal-to-noise (**Supplementary** Fig. 4).

Next, we compared read enrichment surrounding transcriptional start sites (TSS), a commonly used metric for measuring ATAC-seq data quality. We found that concurrent ACCESS-ATAC treatment yielded a slightly higher TSS enrichment than ATAC-seq (13.7x and 12.5x, respectively), while sequential ACCESS-ATAC samples showed substantially decreased TSS enrichment (**Fig. 1E**, **Supplementary** Fig. 4). Decreasing DddSs treatment time during sequential ACCESS-ATAC improved TSS enrichment, although not nearly to the level seen in the concurrent protocol (2.3x for 60-minute, 3.9x for 30-minute, and 7.6 for 15-minute). We conclude that, while concurrent and sequential protocols provide equivalent quality ACCESS data, concurrent ACCESS-ATAC best preserves the accessible chromatin enrichment of ATAC-seq.

We next performed a deeper comparison of K562 concurrent ACCESS-ATAC and ATAC-seq data. We computed the genome-wide correlation between ACCESS-ATAC and ATAC-seq based on read coverage and observed a high Spearman correlation (r = 0.75; **Fig. 1F**). Moreover, the global architecture of accessible peaks is conserved (**Fig. 1G**), suggesting that ACCESS-ATAC does not substantially alter the ATAC signal. At single-nucleotide resolution, DddSs editing events show denser signal than Tn5 insertion frequencies while showing clear visual evidence of TF footprints (**Fig. 1G**). To examine TF footprints more systematically, we produced metaplots centered on ChIP-seq binding events for six TFs. DddSs and Tn5 induce footprints with equivalent foldchange in signal between the peak and trough (**Fig. 1H**), although there are ∼5x more DddSs editing events than there are Tn5 integrations at a given read depth. We observed that DddSs editing footprints are consistently less than half as wide as Tn5 integration footprints (**Fig. 1H**, **Supplementary** Fig. 4), which is presumably due to the considerably smaller size of DddSs (14 kDa) as compared to the Tn5 dimer (106 kDa).

### Transcription factor binding prediction using ACCESS-ATAC

We next assessed whether using the ACCESS signal could improve TF binding site prediction. Accounting for Tn5 motif bias has been shown to be essential for accurate ATAC-seq footprinting analysis^18,19^. To provide a similar correction for DddSs motif bias, we performed DddSs editing followed by whole genome sequencing of purified non-methylated *E. coli* genomic DNA. We obtained >1,000X average coverage across the *E. coli* genome and editing at ∼30% of all cytosines. We found that editing bias was relatively minor and primarily confined to the nucleotides surrounding the target cytosine (**Fig. 1C**). To enable DddSs motif bias correction in ACCESS- ATAC experiments, we trained a convolutional neural network (CNN) model to predict the observed edit counts based on one-hot encoded *E. coli* DNA sequences (**Fig. 2A**). Evaluation on held-out chromatin regions indicated that our model accurately predicted the edit counts for cytosines across the genome (Spearman r = 0.94) and recapitulated DddSs editing profiles at single-nucleotide resolution (**Fig. 2B-C**). We next assessed if this model, trained on *E. coli* DNA, could be transferred to the human genome to account for DddSs bias. We applied the model to predicted DddSs editing counts using the hg38 reference genome and generated the predicted, observed and bias-corrected signal around TF binding site centers (±50bp) identified by ChIP-seq. We found that for many TFs, the observed edit counts around their sequence motifs were strongly influenced by DddSs enzyme bias and a clearer footprint pattern was observed after correcting for bias (**Fig. 2D**; **Supplementary** Fig. 5). These results demonstrate that our CNN model effectively captures DddSs motif bias and enables single-nucleotide resolution bias correction.

**Figure 2:**
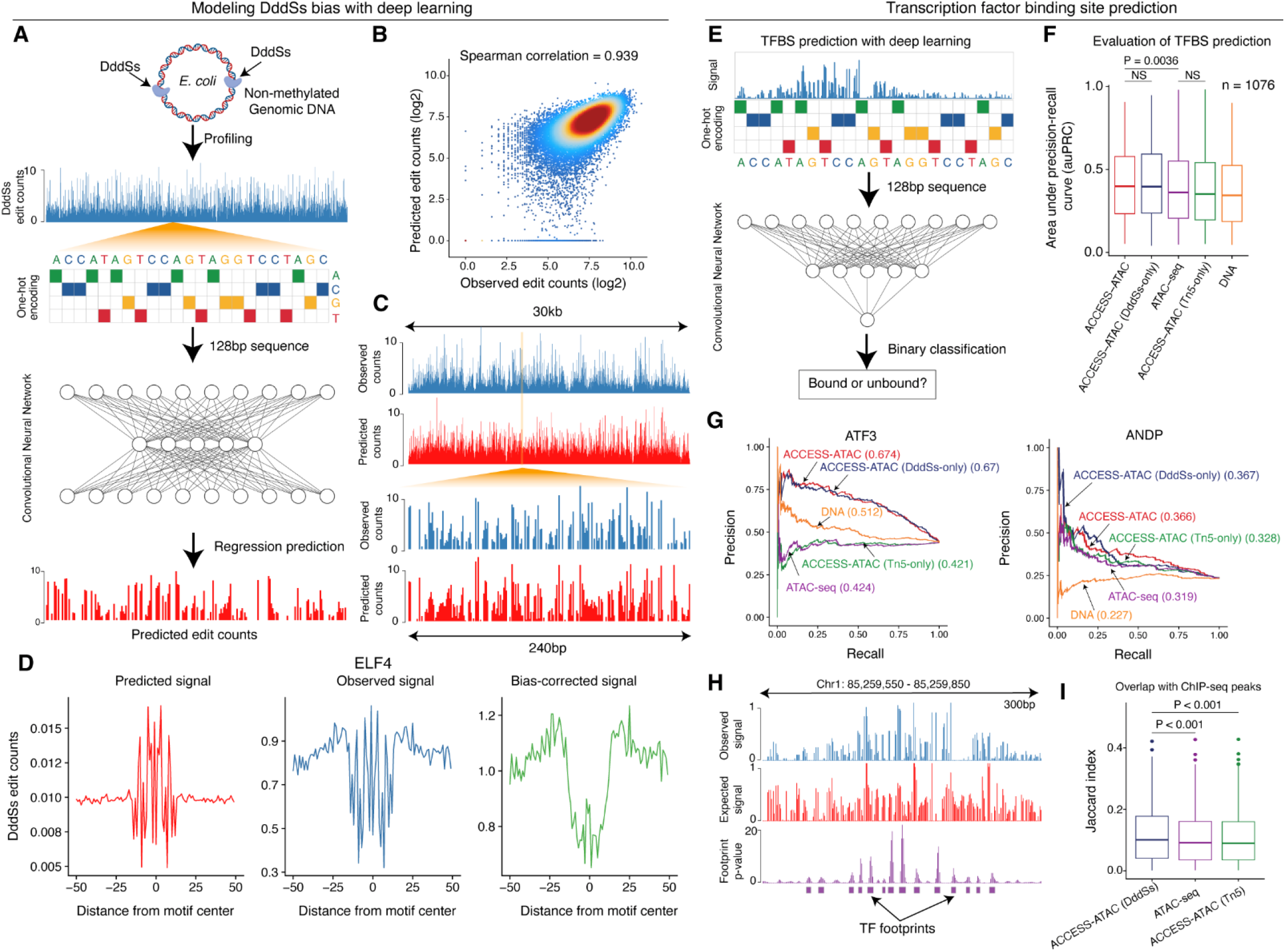
ACCESS-ATAC-seq improves transcription factor binding site prediction. (A) Schematic depicting deep learning-based modeling of DddSs editing bias using high-coverage *E. coli* genomic DNA editing data. (B) Scatter plot showing predicted and observed editing counts for held-out regions (30kb) of the *E. coli* genome. Each dot represents a single nucleotide, and colors refer to density of the nucleotides. (C) Representative tracks showing observed (blue) and predicted (red) DddSs genomic DNA editing counts in a 30-kb stretch (top) and a 240-bp subregion (bottom) of the *E. coli* genome. (D) Single-nucleotide resolution signal for predicted (red), observed (blue) and bias-corrected (green) DddSs edit counts around K562 ELF4 ChIP-seq binding sites from K562 ACCESS-ATAC data. (E) Schematic depicting deep learning-based prediction of transcription factor binding sites using DNA sequence and open chromatin profiles. (F) Evaluation of auPRC comparing TFBS predictions with ChIP-seq binding sites for 1076 K562 motifs using ACCESS-ATAC Tn5 and DddSs signal (red), DddSs only (blue), or Tn5 only (green), ATAC-seq Tn5 signal (purple) or DNA sequence only (orange). Line shows median auPRC, boxes show quartiles, and lines show outliers. (G) Precision-recall curves of TFBS prediction performance for K562 ATF3 and ANDP ChIP-seq binding sites using the indicated tracks. auPRC for each track is reported in parentheses. (H) Representative tracks showing *de novo* footprint detection results using ACCESS-ATAC data in a 300-nt genomic region. Top: observed DddSs edit counts. Middle: predicted DddSs bias using the deep-learning model trained on the *E. coli* genome. Bottom: estimated p-value of TF footprints. (I) Jaccard index of detected footprints using ACCESS-ATAC DddSs signal (blue) or Tn5 signal (green) or ATAC-seq Tn5 signal (purple). For F and I, center line shows median, boxes show quartiles, and lines show outliers.

To determine whether ACCESS-ATAC improves TF binding site prediction, we trained deep learning models using a CNN model similar to the maxATAC approach^27^, which uses DNA sequences and chromatin accessibility signals to predict bound motif instances. To create benchmarking resources, we collected 409 ENCODE TF ChIP-seq datasets in K562 along with 1076 MEME motifs derived from FactorBook^28^. For each motif, we applied the FIMO algorithm^29^ to detect motif-predicted binding sites (MPBSs) based on Position Weight Matrices (PWMs) and overlapped them with corresponding ChIP-seq peaks to obtain the most likely binding sites within peaks to use as true labels. We noticed that, for most TFs, the labels were substantially imbalanced. The fraction of true negatives (MPBS in ACCESS-ATAC peaks but not overlapping ChIP-seq peaks) significantly outnumbered the fraction of true positives, indicating that correctly classifying these labels may pose challenges for predictive models. Next, we trained a CNN-based classifier to predict motif-dependent binding using different input features: i) DNA sequences alone (DNA); ii) DNA plus Tn5 integrations normalized for motif bias from ATAC-seq (ATAC-seq) or ACCESS-ATAC (ACCESS-ATAC (Tn5)) data; iii) DNA plus ACCESS-ATAC DddSs edit events normalized for motif bias (ACCESS-ATAC (DddSs)); or iv) a combination of DNA, DddSs, and Tn5 signal from ACCESS-ATAC (ACCESS-ATAC) (**Fig. 2E**). The results were evaluated on three held-out chromosomes based on the area under the precision-recall curve (auPRC). We found that including DddSs edit signals significantly improved the accuracy of TF binding site prediction (median = 0.398 for ACCESS-ATAC, 0.397 for ACCESS- ATAC (DddSs)), compared with Tn5 (median = 0.36 for ATAC-seq, 0.35 for ACCESS-ATAC (Tn5)) and DNA sequences (median = 0.34; **Fig. 2F-G**; **Supplementary** Fig. 5**; Supplementary Table 1**).

One constraint of this supervised learning approach is that it relies on the existence of corresponding ChIP-seq data from the same context for training models, limiting its applicability in cases where such data is not available. In contrast, TF footprinting algorithms, such as HINT-ATAC^30^ and PRINT^31^, search for footprint-like Tn5 transposase cleavage patterns in ATAC-seq data and subsequently assign TFs to footprints through motif matching. Since we observed that DddSs edit events produce denser signal than Tn5 at single-nucleotide resolution (**Fig. 1G**), we asked if this could also benefit TF footprinting analysis. We implemented a statistical approach, similar to PRINT, to identify genomic regions with significant depletion of DddSs editing counts compared to the background signal (**Fig. 2H**). For a comprehensive comparison, we also applied our method to detect TF footprints using Tn5 insertion patterns derived from conventional ATAC-seq and ACCESS-ATAC data. To quantitatively evaluate the performance of these results, we overlapped the identified TF footprints with peaks from 409 K562 ChIP-seq datasets and computed the Jaccard index to assess their consistency. Notably, our results revealed that TF footprints identified using DddSs editing exhibited significantly higher concordance with the ChIP-seq peaks than those derived from Tn5-based footprinting (**Fig. 2I**, **Supplementary Table 2**). These results underscore the potential of DddSs editing to enhance TF footprinting analysis by providing higher-resolution and more accurate representations of TF binding sites.

### Allele-resolved TF occupancy and co-occupancy imputation from ACCESS-ATAC data

ACCESS-ATAC provides a stencil of bound TFs at a single-allele level, similar to SMF^32^ and Fiber-seq^12^. Because, in contrast to these other methods, ACCESS-ATAC is compatible with short-read sequencing and accessible chromatin enrichment, we posited that ACCESS-ATAC could permit high-coverage single-molecule profiling of TF occupancy across the accessible human genome. We used Ultima Genomics sequencing^33^ to generate high-coverage ACCESS-ATAC datasets in HepG2 and K562 cells to leverage the wealth of TF binding data in these cell lines^34,35^. After alignment and deduplication, we retained 330 million HepG2 and 612 million K562 reads.

We developed the Occupancy Pattern Inference by Editing (OccuPIE) model to impute TF occupancy status at individual alleles from ACCESS-ATAC data using similar principles to a prior approach based on SMF data^32^ (**Fig. 3A**). OccuPIE assumes that a candidate TF binding site can exist in three states: actively occupied by the TF (bound); unoccupied but with accessible chromatin; and unoccupied with inaccessible chromatin. Using 98 HepG2 and 106 K562 ChIP-seq datasets, we used the composite editing patterns surrounding ChIP-seq-bound motif instances to define a TF-specific peak segmentation with three nucleotide ranges, one for the central footprint and two for the flanking regions. We then created a training set by assigning ACCESS-ATAC reads that spanned a ChIP-seq-bound motif into one of the three states through defining thresholds of editing in the footprint and peak regions. Bound reads were defined as having high editing in flanking regions and low footprint editing, unbound, accessible reads as having high editing in flanks and footprint, and unbound, inaccessible reads as having low editing in flanks and footprint. We used these assignments to train a deep learning model to output the occupancy probability for every motif-overlapping allele in our ACCESS-ATAC datasets.

**Figure 3:**
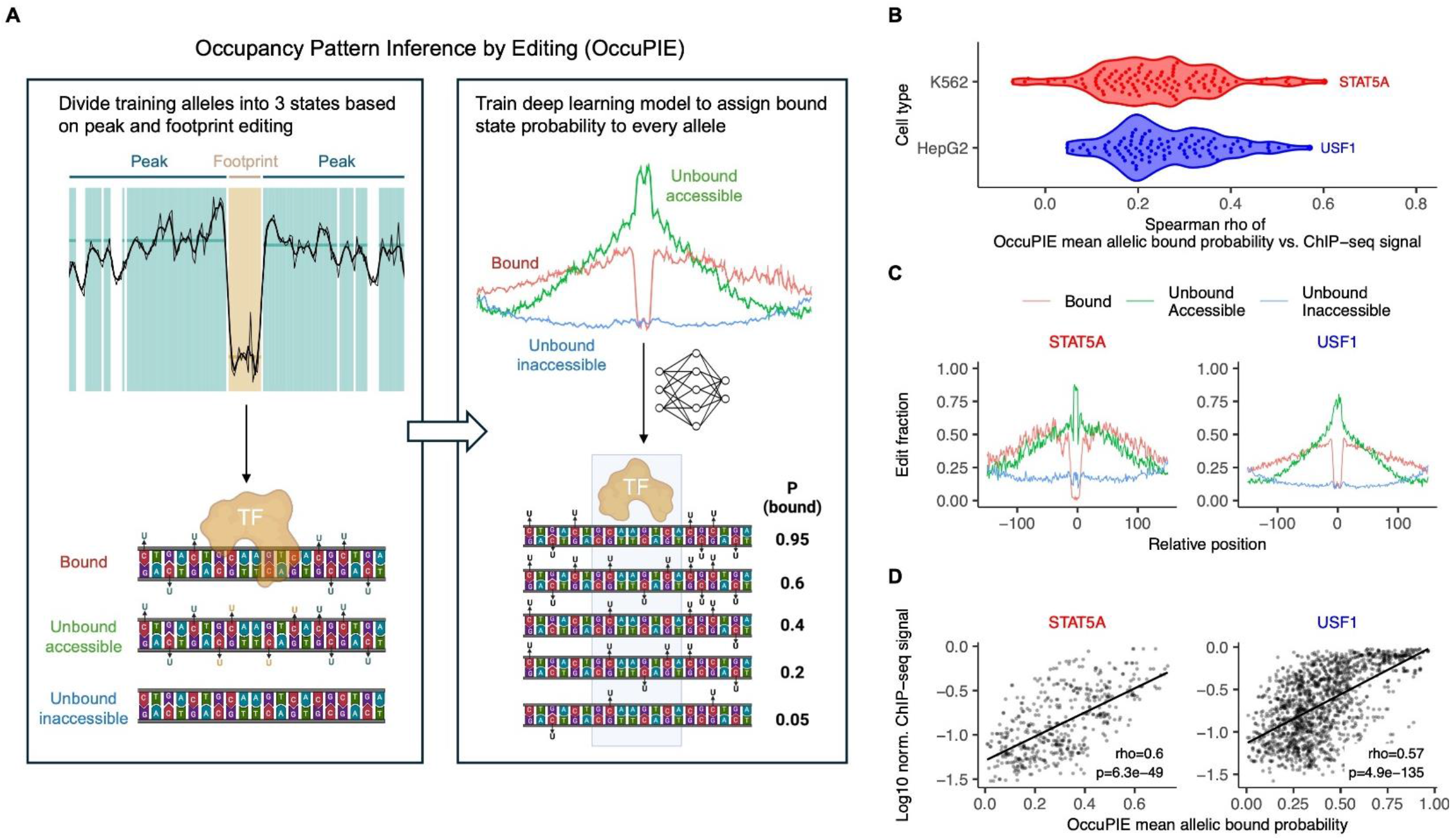
Imputing allele-resolved transcription factor occupancy from ACCESS-ATAC data. (A) Schematic of Occupancy Pattern Inference by Editing (OccuPIE), a model to impute allelic TF occupancy from ACCESS-ATAC data. (B) Evaluation of Spearman correlation between OccuPIE-imputed median allelic occupancy and ChIP-seq read abundance for 98 HepG2 and 106 K562 TFs (C) Edit fractions for reads surrounding K562 STAT5A (left) and HepG2 USF1 (right) ChIP-seq-bound motifs. Reads are separated into bound (red), unbound, accessible (green), and unbound, inaccessible (blue) states according to maximum OccuPIE-assigned probability. (D) Evaluation of mean OccuPIE-imputed bound probability (x-axis) and ChIP-seq read abundance (y-axis) at K562 STAT5A (left) and HepG2 USF1 (right) ChIP-seq bound loci.

To assess whether the output of OccuPIE accurately reflects TF binding propensity, we compared the median imputed allelic occupancy probability at a given motif instance with the abundance of ChIP-seq reads at that locus (ChIP-seq score). For 87 of the HepG2 and 85 of the K562 TFs, there was a significant association between OccuPIE-imputed allelic occupancy and ChIP-seq score across bound loci (Bonferroni-corrected p-value <0.01, **Fig. 3B**, **Supplementary Table 3**). This association was strongest for TFs with wide, robust footprints such as STAT5A and USF1 (**Fig. 3B-D**, **Supplementary** Fig. 6), indicating that TFs that strongly impede editing when bound allow for the most robust occupancy imputation. We found that TFs show a wide dynamic range in their allelic occupancy. For example, HepG2 USF1 loci ranged from 1-97% occupancy, with a median of 36% of bound alleles (**Fig. 3D**). OccuPIE-imputed occupancy showed consistently improved association with ChIP-seq score as compared with the predefined training labels (median 19% increased Spearman correlation, Supplementary Fig. 6), validating its utility.

We next used OccuPIE to assess co-occupancy patterns among 64 TFs in HepG2 and 64 TFs in K562 with robustly imputed allelic occupancy (Spearman r > 0.2 with ChIP-seq score). We examined ChIP-seq-bound motif pairs within 100 nt, removing motif pairs for which the central footprints for the two TFs have overlapping nucleotides, as OccuPIE cannot independently assess occupancy for such pairs. Altogether, we assessed co-occupancy at a total of 658,230 adjacent motif pairs in HepG2 and 1,105,054 in K562. To control for factors such as motif strength that might cause a TF to bind more or less frequently at a given motif instance, we calculated an “expected” co-occupancy for every adjacent motif pair under the baseline assumption that binding occurs independently (Mean P(boundTF1) * Mean P(boundTF2)). For each motif pair, we then compared the observed co-occupancy for all ACCESS-ATAC alleles overlapping both motifs with this baseline value to determine whether the two TFs co-occupy more often than expected (observed – expected co-occupancy, **Fig. 4A**).

**Figure 4:**
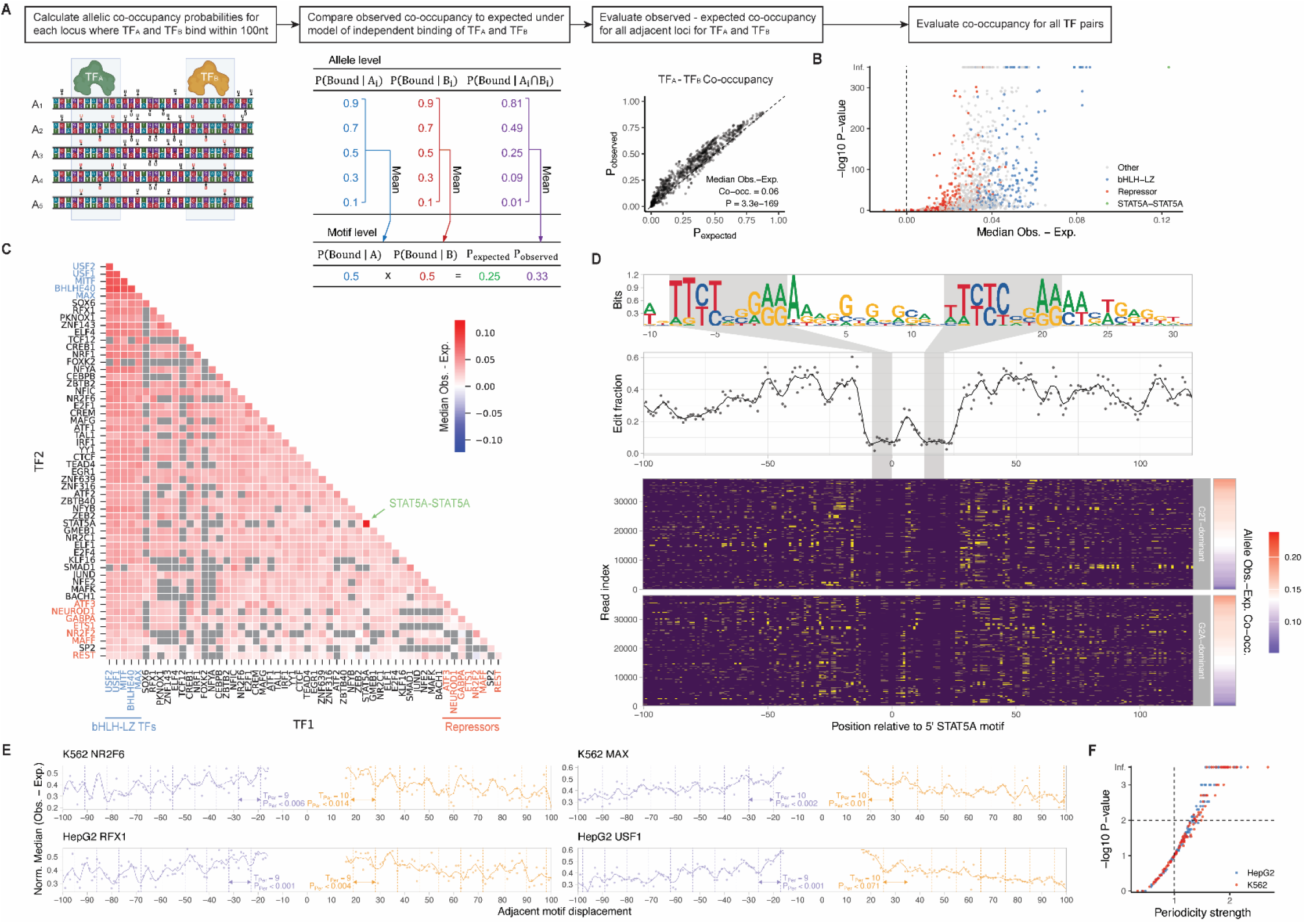
Transcription factors show enriched co-occupancy of adjacent motifs with helical periodicity. (A) Schematic showing the procedure used to assess allelic co-occupancy enrichment. (B) Evaluation of magnitude (x-axis, median observed – expected co-occupancy) and statistical significance (y-axis, -log10(p)) of allelic co-occupancy for 2,373 TF pairs in K562. (C) Heat map showing median allelic co-occupancy enrichment for all 2,373 TF pairs in K562. In B-C, TF pairs that include a group of bHLH-LZ TFs with enriched co-occupancy (blue), a set of repressor TFs with minimally enriched co-occupancy (red), and STAT5A-STAT5A (green) are highlighted. (D) Analysis of K562 ACCESS-ATAC editing in 67,951 alleles at 460 K562 STAT5A ChIP-seq- bound loci with fixed 21-nt distance between motif centers. Top: motif logo showing positional nucleotide representation at the 460 loci. The STAT5A primary motifs are highlighted. Middle: Positional DddSs edit fractions for alleles at the 460 loci. STAT5A motif positions are highlighted, and a smoothed trendline is included. Bottom: Allelic editing patterns for all 67,951 reads at the 460 loci. Unedited (purple) and edited (yellow) positions are shown. Alleles are separated by C2T-dominant and G2A-dominant editing patterns and sorted by observed – expected allelic co-occupancy. (E) Normalized median position-resolved observed – expected co-occupancy for all loci for which the listed TF is within 100-nt of any of the 64 analyzed TFs. The motif for the listed TF is always centered at 0 on the x-axis. The dominant period (Tp) and Bonferroni-corrected p-value from fast Fourier transform (FFT) analysis are shown for motifs upstream (red) and downstream (blue) of the listed TF, and smoothed trendlines are included. (F) Evaluation of periodicity strength and Bonferroni-corrected p-value derived from FFT analysis of median position-resolved observed – expected co-occupancy measurements for 64 K562 (blue) and HepG2 (red) TFs. For each TF, all loci bound by any other TF within 100-nt are included, analysis is performed separately for motifs upstream and downstream, and values are reported for the dominant period. (Fig5F_co-occupancy volcano). (G) Schematic summarizing the findings from periodicity analysis. For over half of analyzed TFs, co-occupancy is enriched with periodicity consistent with the 10.5-nt helical turn of DNA. We posit that these TFs are more prone to interact either directly with each other or with a common cofactor when bound in major grooves positioned on the same face of the DNA helix.

Our analysis supports the conclusion that TFs tend to co-occupy adjacent binding sites more frequently than would be expected by chance. We observe significantly enriched co-occupancy for >85% of TF pairs in HepG2 and K562 (Bonferroni-corrected p < 0.01, **Fig. 4B**, **Supplementary** Fig. 7, **Supplementary Table 4**). TFs of the bHLH-LZ class such as USF1, MITF, and MAX show enriched co-occupancy with the majority of other TFs and especially strong co-occupancy with each other (**Fig. 4C**, **Supplementary** Fig. 7). While these factors are known to homo- and heterodimerize^36^, the inter-motif distances explored in this analysis are larger than can easily be explained by dimerization. TFs that function as repressors such as REST, MAFF, and NR2C2 show the lowest co-occupancy enrichment.

Of all analyzed adjacently bound TF pairs in K562, homotypic STAT5A-STAT5A pairs showed the most robustly enriched co-occupancy (**Fig. 4C**). This enrichment was primarily driven by over 500 loci with ChIP-seq-bound STAT5A motif pairs for which the motif centers are separated by 20-25 nt (**Supplementary** Fig. 7). While these loci exhibit a wide range of STAT5A occupancy levels (median allelic occupancy probabilities from 10-75%), they show uniformly highly enriched allelic co-occupancy (**Fig. 4D**, **Supplementary** Fig. 7). STAT5A binds as a homodimer to a palindromic TTCNNNGAA consensus, and Stat factors including STAT5A are known to tetramerize upon persistent activating stimuli, enabling binding to lower affinity sites^37–39^. K562 cells possess BCR/ABL translocation, a known driver of Stat tetramerization^40^. Our allele-resolved analysis shows that, at loci with properly positioned motif pairs, activated STAT5A in K562 cells interacts with DNA predominantly as a tetrameric unit in all-or-nothing fashion.

We further assessed whether co-occupancy is dependent on the distance between the two bound motifs. To maximize power for this analysis, we assessed spatially resolved co-occupancy enrichment for each TF when paired with any adjacent TF. Intriguingly, we found that, in addition to a steadily decreasing propensity for co-occupancy at increasing distances, certain TFs show enriched co-occupancy at periodic intervals (**Fig. 4E**, **Supplementary** Fig. 7). Fast Fourier transform (FFT) analysis revealed that co-occupancy enrichment shows significant periodicity for 39 HepG2 and 32 K562 TFs in range of the 10.5 bp periodicity of the DNA helical turn (**Fig. 4F**, **Supplementary** Fig. 7, **Supplementary Table 5**). TFs with periodic co-occupancy span multiple families^1^, suggesting that this property is not dependent on specific DNA-binding or protein-protein interaction domains. Periodic co-occupancy was often evident throughout the entire +/- 100-nt evaluated range, suggesting that this trend persists across a large number of helical turns of the DNA. We were unable to assess periodicity between specific TF pairs due to data sparsity. Altogether, our analysis reveals that many TFs show periodic co-occupancy on individual DNA molecules.

### Single-cell ACCESS-ATAC

Because the resolution of single cell ATAC-seq (scATAC-seq) is inherently limited, we next asked whether we could combine ACCESS with scATAC-seq (scACCESS-ATAC) to generate high-resolution chromatin accessibility profiles at the single-cell level (**Fig. 5A**). We performed a proof-of-principle experiment in which we performed arrayed CRISPR-KO of 11 chromatin-associated proteins as well as a non-targeting control gRNA in HepG2 and K562 cells. We also collected two additional control datasets by performing scATAC and scACCESS-ATAC on these two cell lines without gRNA treatment. We adapted the 10X Chromium GEM Single Cell ATAC v2 protocol, adding DddSs treatment and DddSs-I quenching prior to treatment with barcoded Tn5 (sequential ACCESS-ATAC) and replacing the polymerase mix in the two PCR steps (in-droplet and after purification) with Q5U. The cells were assigned to the CRISPR-KO treatment they received based on the combination of cellular and Tn5 barcode. Using super-loading, we recovered a total of 30,457 cells with an average of 2,387 unique fragments and TSS enrichment score of 8.3 per cell after controlling the data quality (**Supplementary** Fig. 8). These results indicate that DddSs treatment is compatible with the 10x scATAC-seq protocol.

**Figure 5:**
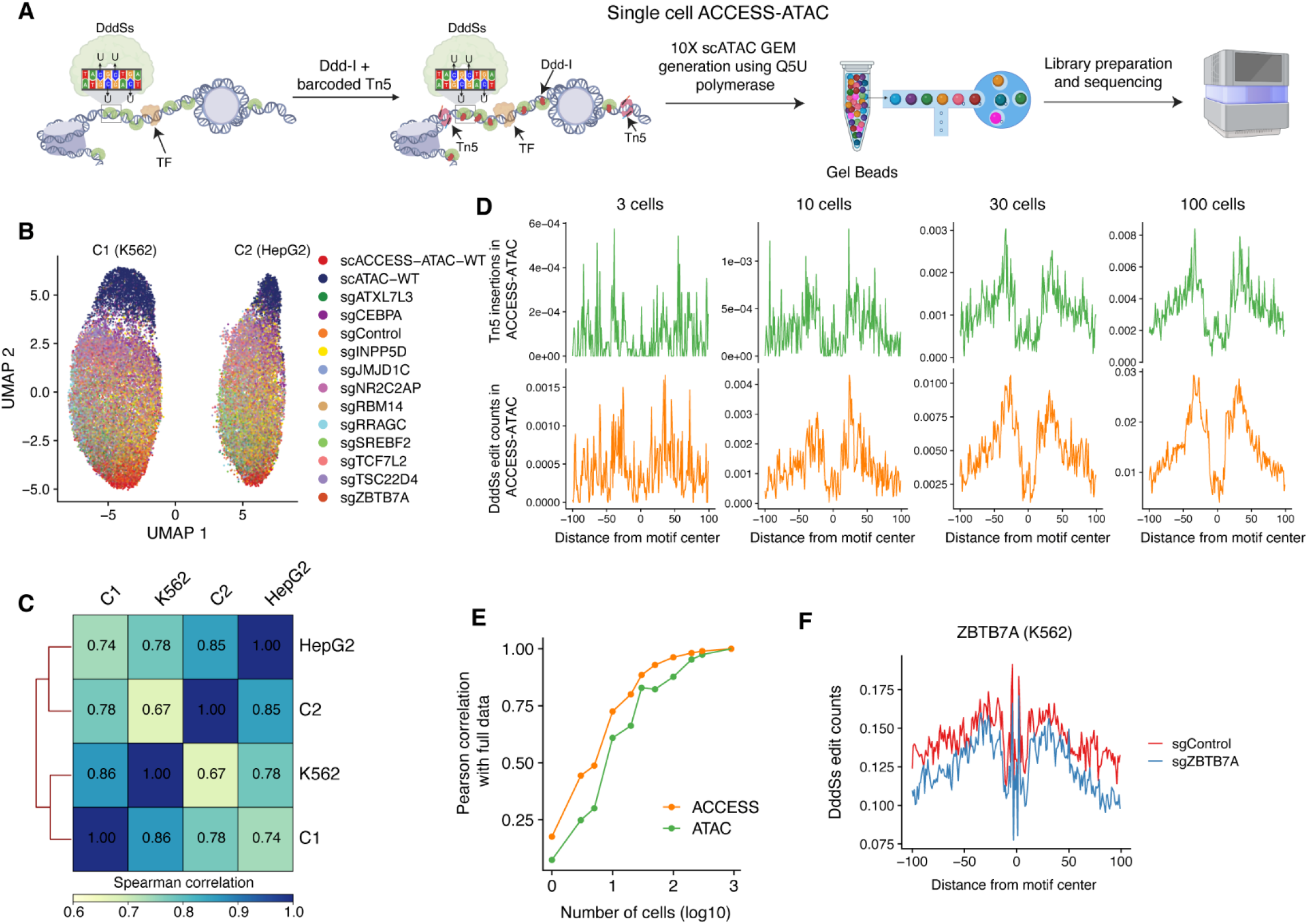
ACCESS-ATAC improves resolution of single-cell chromatin accessibility profiling. (A) Schematic of single-cell ACCESS-ATAC. Sequential treatment with DddSs, DddSs inhibitor, and Tn5 are used prior to encapsulation of nuclei following a minimally altered 10X scATAC workflow. (B) Barplot showing the number of valid cells after filtering by quality for each indexed sample within the scACCESS-ATAC experiment. (C) UMAP embedding of integrated scACCESS-ATAC data as colored by clusters. (D) Heatmap showing the genome-wide Spearman correlation among the two scACCESS-ATAC clusters, and K562 and HepG2 bulk ATAC-seq datasets. (E) Metaplots showing Tn5 insertions and DddSs editing at CTCF binding sites in K562 for different number of cells. (F) UMAP plot showing the scACCESS-ATAC data as colored by sgControl and sgZBTB7A treatments. (G) Metaplots showing DddSs editing at ChIP-seq-bound ZBTB7A motifs in pseudo- bulk sgControl (red) and sgZBTB7A (blue) populations assigned as K562 (left) or HepG2 (right).

We next assessed whether HepG2 and K562 cells could be distinguished using scACCESS-ATAC data. To do this, we aggregated the cells from all experiments and identified 94,366 peaks based on which a cell by peak count matrix was constructed using the ATAC profiles. We reduced the dimensionality using Latent Sematic Indexing (LSI) and projected the cells into a 2-dimensional space using Uniform Manifold Approximation and Projection (UMAP). We observed that different CRISPR-KO treatments were separated from the WT and enriched in the different areas in the UMAP embedding space, indicating that the KO of these 11 proteins differently altered the chromatin accessibility and that scACCESS-ATAC was able to capture these differences (**Fig. 5B**). We next clustered the cells and identified two major populations, designated as C1 and C2, which presumably correspond to the two component cell lines (**Fig. 5B**). To annotate the clusters, we generated the pseudo-bulk ATAC profiles for C1 and C2 and compared them to the ATAC-seq data for K562 and HepG2 cells obtained from ENCODE project. We observed a strong correlation between C1 and K562 (Spearman r = 0.86) and between C2 and HepG2 (Spearman r = 0.85) (**Fig. 5C**). We therefore annotated C1 as K562, and C2 as HepG2. Together, these results demonstrate that scACCESS-ATAC can effectively distinguish between different cell states and types within complex cellular mixtures.

Single-cell ATAC-seq has been used to compare TF binding activity in pseudo-bulk populations. To evaluate whether the increased resolution of scACCESS-ATAC could improve this application, especially for rare populations, we performed a proof-of-concept analysis. We subsampled K562 cells and assessed CTCF binding by plotting Tn5 insertions or DddSs edit counts surrounding K562 CTCF ChIP-seq-bound motifs for different number of cells (**Fig. 5D**). As expected, we observed that the clarity of the peak and footprint profile improved as increasing cell numbers were included. Moreover, using DddSs editing signal consistently produced a cleaner profile than using Tn5 insertion signal at a given cell number. To quantify this improvement, we calculated the Pearson correlation between the subsampled CTCF profile and the profile obtained by using all cells (**Fig. 5E**). We found that DddSs editing data requires 2-3-fold fewer cells than Tn5 insertion data to achieve a similar quality profile (similar Pearson correlation) across a wide range of subsampled inputs, demonstrating the superior performance of scACCESS-ATAC for detecting TF binding activity even with limited cell numbers. We next investigated the impact of CRISPR-KO on TF binding activity. We created a pseudo-bulk profile for each cell type and experimental condition by aggregating the scACCESS-ATAC data. Next, we performed differential footprinting analysis by comparing the ACCESS profiles between sgControl and sgZBTB7A based on the ChIP-seq peaks of ZBTB7A in K562. We observed that sgZBTB7A treatment significantly diminished ZBTB7A binding (**Fig. 5F**). These findings demonstrate the utility of the additional information recoverable from the single-cell level editing patterns provided by scACCESS-ATAC in interrogating TF binding differences across cellular contexts.

## Discussion

In this work, we establish ACCESS-ATAC, an approach that leverages dsDNA cytosine deamination to increase the resolution of chromatin accessibility measurement across the accessible genome. Recently developed techniques such as Fiber-seq^12^, SMAC-seq^13^, NOMe-seq^15^, and SMF^14^ have demonstrated that enzymes that mark accessible nucleotides without terminating NGS reads allow for insight into allele-level chromatin architecture and TF binding. A recent preprint also reports the use of DddSs (called SsDddA) in conjunction with long-read sequencing^25^. However, none of these prior techniques has been shown to be compatible with ATAC-seq or with high-throughput single-cell profiling. We show that ACCESS-ATAC retains equivalent accessible chromatin enrichment as ATAC-seq and is compatible with single-cell chromatin accessibility profiling. Thus, ACCESS- ATAC provides an easily implemented, high-resolution approach to profile bulk and single-cell chromatin accessibility and allelic TF occupancy, and we have developed computational pipelines to normalize and analyze the resulting data. We note that a method similar to sequential ACCESS-ATAC has been reported using a distinct Ddd ortholog^41^, and it will be informative to compare these methods in the future, especially since our data suggest concurrent enzyme treatment provides optimal performance.

We show that ACCESS-ATAC outperforms ATAC-seq at identifying TF binding sites using two complementary analyses: supervised deep learning-based prediction and *de novo* TF footprint detection. This is at least partly due to information content—each paired-end ACCESS-ATAC read contains dozens of editable nucleotides and >10 edits on average as compared with only two Tn5 integration events. Moreover, ACCESS footprints are considerably narrower than those generated by ATAC cleavage and thus align more precisely with underlying motifs. It is possible that disambiguation of footprints in loci with multiple adjacent binding events contributes to improved TFBS prediction. The TFBS prediction method used in this work sums positional DddSs edit counts, but taking full advantage of ACCESS signal by exploiting allelic editing information could be a promising approach to improve TFBS prediction.

In addition to improving the resolution of population-level TFBS prediction, ACCESS-ATAC allows for allele-level TF occupancy imputation. We build a model, OccuPIE, that uses DddSs editing in peak and footprint regions to classify TF occupancy on individual alleles. We find association between OccuPIE-imputed allelic occupancy and ChIP-seq read depth for over 100 TFs, but the strength of this association varies. The variable strength of this association likely depends on both the robustness of ACCESS footprints and ChIP-seq data. TF footprints vary in their width and strength due to TF-intrinsic properties, and ACCESS signal also depends on the GC content of the footprint region due to use of a cytosine deaminase. One current limitation of ACCESS-ATAC data is that individual reads carry edits either from the top (C-->T) or bottom (G-->A) strand. Resolving edits on both strands would be expected to improve allelic occupancy imputation. Beyond improving occupancy imputation, it is also worth noting that ChIP-seq read depth may be an imperfect gold standard because adjacent binding site presence, PCR bias, and other experimental artifacts can affect read abundance at a locus.

The ability of ACCESS-ATAC to measure allelic TF occupancy allows assessment of allelic co-occupancy for thousands of TF pairs across over a million adjacent bound motifs. We find that >85% of TF pairs co-occupy adjacent binding sites more often than expected under an independent co-occupancy model. This finding is in line with prior analysis of SMF data in mESCs^32^, although our analysis is over an order of magnitude more comprehensive. This widespread co-occupancy enrichment across TF families is consistent with models of chromatin accessibility, which suggest that TFs interact through a non-specific cooperative logic to outcompete nucleosome binding of DNA^42,43^.

We find that TF co-occupancy preference shows two spatial major trends. First, co-occupancy decays as the distance between motifs increases. Second, we find that over half of analyzed TFs show periodicity in the degree of co-occupancy enrichment approximating the DNA helical turn. Helical periodicity has been observed in the preferences of TFs to bind to nucleosomal DNA^44,45^. Additionally, several individual TFs have been observed to show oscillating co-binding preference^46–49^. Our work reveals that such periodic co-occupancy preference is widespread across TFs and occurs at the level of binding to individual DNA molecules. The vast majority of TFs bind to the major groove of DNA^50^, and major grooves are positioned on the same plane of the DNA at ∼10.5 nt intervals. Thus, the most likely explanation for the observed periodic co-occupancy is that positioning TFs on the same plane maximizes the ability of TFs to interact with each other or with common cofactors^51^. The detection of periodic co-occupancy for TFs separated by up to 100 nucleotides favors a model of shared interaction with a multivalent binding partner such as the Mediator complex^52^, although further work is required to understand the driving mechanism(s). It has also been shown that protein occupancy in one major groove can promote binding at the adjacent major groove by altering DNA vibration^53^, thereby altering the width of the adjacent major groove. It is unclear if this effect could extend to the distances observed in this work, but it could provide an additional mechanism to explain periodically enriched co-occupancy.

It is worth noting that our quantification of allelic co-occupancy is likely to be overly conservative. To control for locus-specific parameters such as motif strength, we measure the increased co-occupancy of a TF pair compared to an independent model based on the allelic occupancy of each TF at that locus. However, cooperative TF interactions would be expected to increase allelic occupancy of each TF on top of allelic co-occupancy, weakening the apparent enrichment that we measure. We also only consider motifs with evidence of binding from ChIP-seq, so apparent co-occupancy enrichment may be further weakened by the fact that adjacent motifs that do not benefit from each other’s binding may lack ChIP-seq signal and thus not be considered. Generative computational models of chromatin accessibility and TF binding^46,54^ and perturbational approaches to assessing accessibility and TF binding^49,55^ are well-positioned to dissect the relative impact of TF-TF distance and periodicity as compared to primary motif features.

Altogether, ACCESS-ATAC provides a unique solution for high-resolution, single-cell and single-molecule genome-wide chromatin accessibility profiling.

## Acknowledgments

The authors thank Grigoriy Losyev, Sitara Roy, and Mandovi Chaterjee for technical assistance, Joseph Mougous for reagents, and Ultima Genomics for sequencing.

## Author contributions

Conceptualization: E.G., R.I.S.; Experimental investigation: E.G., R.I., M.J.F., Q.v.P., J.R.; Computational analysis: T.Y., Z.L., J.Z., Q.v.P.; Funding acquisition and project supervision: P.A.C., L.P., R.I.S.; Writing: T.Y., Z.L., E.G., R.I., M.J.F., Q.v.P., J.Z., J.R., P.A.C., L.P., R.I.S.

## Data and materials availability

Data and materials will be made available upon publication. Code is available at https://github.com/pinellolab/ACCESS-ATAC-seq.

## List of Supplementary Materials

Materials and Methods

Supplementary Text

Figs. S1 to S8

Tables S1 to S6

