## Supplement for "Deaminase-mediated chromatin accessibility profiling with single-allele resolution"

### Materials and Methods

#### Expression of Ddd enzymes through *in vitro* transcription and translation

Amino acid sequences of six Ddd enzymes were obtained from published work. Nucleotide sequence was codon optimized for *E. coli* expression using IDT Codon Optimization tool, and open reading frames containing T7 promoter and terminator stubs (Supplementary Text) were ordered from Twist Bioscience. T7 promoter and terminator extension primers (Supplementary Text) were used to PCR amplify each Ddd enzyme, and PCR purified products were used as input for *in vitro* transcription and translation using the NEB PURExpress® In Vitro Protein Synthesis kit. Activity of bulk protein obtained from *in vitro* transcription and translation was titrated using the *in vitro* dsDNA deamination assay, and concentrations of each Ddd enzyme that yielded ~50% *in vitro* dsDNA deamination were used in ACCESS-WGS.

#### Bacterial expression and purification of Ddd enzymes

A plasmid allowing for dual bacterial expression of DddA and DddA-I (pETDuet_DddA-DddA-I) was a kind gift from Joseph Mougous. We subcloned pETDuet_DddSs-DddSs-I by ordering *E. coli* codon optimized sequences for this enzyme (adding a His tag) and inhibitor based on Genbank sequences. The sequence of these inserts is provided in Supplementary Text. The enzymes and inhibitors were purified based on a published protocol[56]. The plasmids were transformed into BL21 Codon Plus cells for recombinant protein expression. The cells were cultured in LB/Ampicillin medium to an optimal cell density at $OD_{600} = 0.6$ and induced at 18°C overnight. The cells were harvested, resuspended in lysis buffer (50 mM Tris pH 8.0, 500 mM NaCl, 10 mM imidazole, 1 mM TCEP, 1mM PMSF, 1 tablet of protease inhibitor) and lysed by passing through a french press three times. After centrifugation at 27,000g for 40 min to remove debris, the lysate was passed through 4 mL of Ni-NTA resin (MCLAB Catalog #NINTA-100) three times. The resin was washed with 12 column volumes (CV) of wash buffer 1 (50 mM Tris pH 8.0, 500 mM NaCl, 10 mM imidazole, 1 mM TCEP) and 5 CV of wash buffer 2 (50 mM Tris pH 8.0, 500 mM NaCl, 20 mM imidazole, 1 mM TCEP). The deaminase-inhibitor complex was eluted using elution buffer (50 mM Tris pH 8.0, 500 mM NaCl, 300 mM imidazole, 1 mM TCEP) and dialyzed in 8 M urea buffer (50 mM Tris pH 7.5, 500 mM NaCl, 10 mM imidazole, 1 mM TCEP, 8 M urea) overnight. The dialyzed sample was passed through Ni-NTA resin three times and washed with 12 CV of 8 M urea buffer.

For Deaminase (DddA/DddSs) Isolation: The Ni-NTA resin was washed sequentially with 6 CV of 6 M, 4 M, 2 M, and 1 M urea buffer. After a final wash with 12 CV of 0 M urea buffer, the deaminase was eluted using elution buffer. *For Deaminase Inhibitor (DddA-I/DddSs-I) Isolation:* The flowthrough from the Ni-NTA purification post 8 M urea dialysis and the 8 M urea wash fractions were pooled. The pooled fractions underwent step-dialysis using 6 M, 4 M, 2 M, 1 M, and 0 M urea buffer.

The deaminase and deaminase inhibitor samples were each dialyzed in 0 M urea buffer overnight and concentrated. The proteins were further purified on a Superdex 75 10/300 GL column (Cytiva, Catalog #17517401) with a mobile phase of 20 mM Tris pH 7.5, 200 mM NaCl, 2 mM DTT, 5% glycerol, flash frozen, and stored at -80°C until use. All purified proteins were characterized by SDS-PAGE and ESI-MS (Q Exactive, Thermo Scientific). Data analysis of the ESI-MS spectra was performed using UniDec software[57].

#### *In vitro* dsDNA deamination assay

Deamination of double-stranded DNA (dsDNA) by Ddd enzymes was measured through a DpnII restriction digest assay. A brief summary follows, and a full protocol is provided (Supplementary Text). Two complementary 90-nt oligos were ordered from IDT with a central GATC sequence surrounded by A:T basepairs (Supplementary Text).

These oligos were annealed at 25 uM in IDT Nuclease Free Duplex Buffer through heating to 95 deg C followed by gradual cooling to 25 deg C. Deamination was performed in a 10 uL reaction with final concentrations of 20 mM Tris-HCl pH 7.4, 200 mM NaCl, 1 mM DTT, 1 uM dsDNA substrate. Deamination was detected by DpnII (NEB) restriction digest. Deamination efficiency was measured using D1000 Tapestation (Agilent).

**ACCESS followed by whole genome sequencing (ACCESS-WGS)**

A full protocol for ACCESS-WGS is provided in Supplementary Text. A brief summary follows. HCT116 cells were harvested by trypsinization, and 2.5*10^5 cells per reaction were lysed for 3 minutes on ice followed by dilution and centrifugation to recover intact nuclei. Nuclei were then treated for 30 min at 37 deg C in a 25 uL reaction with a concentration of Ddd enzyme titrated to have ~50% deamination activity in the dsDNA deamination assay. Treated genomic DNA was purified using the Zymo Genomic DNA clean and concentrator 25 kit. Tagmentation was performed by 5 minute incubation at 55 deg C of 100 ng of ACCESS-treated genomic DNA in a 50 uL reaction using 0.5 uL of Tagmentase (Tn5 transposase) - loaded (Diagenode). Tagmented genomic DNA was purified using the Zymo DNA clean and concentrator 5 kit. Nextgen sequencing library prep was then performed using 2x Q5U mastermix (NEB).

**ACCESS-ATAC**

Full protocols for ACCESS-ATAC concurrent and sequential treatment are provided in Supplementary Text. A brief summary follows. K562 or HepG2 cells were harvested, and 2.5*10^5 cells per reaction were lysed for 3 minutes on ice followed by dilution and centrifugation to recover intact nuclei. For concurrent ACCESS-ATAC, nuclei were treated for 30 min at 37 deg C in a 25 uL reaction with 250 nM DddSs and 1.25 uL of Tagmentase (Tn5 transposase) - loaded (Diagenode). For sequential ACCESS-ATAC, nuclei were treated for 15-30 min at 37 deg C in a 25 uL reaction with 250 nM DddSs, then DddSs activity was quenched by treatment for 2 min at 37 deg C with 5-10-fold molar excess of DddSs-I. Then nuclei were treated for an additional 30 min at 37 deg C with 1.25 uL of Tagmentase (Tn5 transposase) - loaded (Diagenode). ACCESS-ATAC-treated DNA was purified using the Zymo DNA clean and concentrator 5 kit. Nextgen sequencing library prep was then performed using 2x Q5U mastermix (NEB). Nextgen sequencing was performed using Element AVITI at Quintara Biosciences or Ultima Genomics UG100 at Ultima as specified.

**Single-cell ACCESS-ATAC**

K562 lentiviral CRISPR-Cas9 knockout lines were made for 12 gRNAs, each cloned into lentiCRISPRv2-FE PuroR (Addgene 186746). gRNA sequences are listed in **Supplementary Table 6**. Lentiviral production and Puromycin selection were performed as described.

On the day of the experiment, 2.0*10^5 of each cell gRNA-treated cell population, as well as HepG2-wt and K562-wt cells, were lysed, and the equivalent gRNA-treated populations of K562 and HepG2 nuclei were combined. Nuclei were treated as in the sequential ACCESS-ATAC protocol with 15-minute DddSs treatment time up through the DddSs-I quenching step. Barcoded Tn5 treatment was then performed using the Scale Biosciences scATAC Pre-Indexing Kit, with 30-minute Tn5 treatment. All nuclei were combined and processed as specified in the 10X Chromium Next GEM Single Cell ATAC Kit v2 with the following changes from the standard protocol. 50,500 nuclei were loaded into the GEM generation step. Barcoding enzyme mix was replaced with 2 uL Q5U Hotstart DNA polymerase (NEB) and 0.5 uL ET SSB (NEB) in step 2.1. Eight linear amplification cycles were used in step 2.5. In step 4.1, Single Index N Set A primer was replaced with Scale Bio is700 primer, and 2x NEBNext Q5U mastermix was used in place of Amp Mix. Nextgen sequencing was performed using Element AVITI at Quintara Biosciences.

**Processing of ACCESS-ATAC-seq data**

To process the edit-rich ACCESS-ATAC-seq datasets, we employed an iterative alignment approach using Bowtie2[24] (v2.5.3) parameter. In the first iteration, reads were aligned with the "--very-sensitive" preset and default maximum mismatch penalty ("--mp") of 6. Unmapped reads from each iteration were subsequently re-aligned in further iterations, reducing the mismatch penalty by 2 in each step. This process included a total of five iterations, and alignments from all iterations were merged into a single dataset. Deduplication of aligned reads was performed using czid-dedup (v0.1.2). Multi-mapped reads were then removed with a custom Python script, retaining only uniquely aligned reads from chromosomes 1-22 and X.

**ACCESS-ATAC-seq quality control metrics**

We established three quality control modules for ACCESS-ATAC-seq: transcription start site enrichment (TSS Enrichment), accessible chromatin log fold change (AC-LFC), and peak-trough log fold change (PT-LFC). For TSS Enrichment, we adapted the analysis script from the ENCODE ATAC-seq pipeline (github.com/ENCODE-DCC/atac-seq-pipeline/). AC-LFC and PT-LFC, on the other hand, are based on the editing activity of DddSs. To calculate AC-LFC, we utilized an in-house Python script to bin the genome by accessibility scores and determine the fraction of edited bases in ACCESS-ATAC-seq reads mapped to each accessibility bin. Chromatin accessibility scores for HepG2 (ENCFF262URW), K562 (ENCFF600FDO) and HCT116 (ENCFF962GFP) cell lines were obtained from existing ATAC-seq experiments in the ENCODE Project. AC-LFC was defined as the log fold change between the edit fraction in low-accessibility regions (accessibility<0.1) and high-accessibility regions (accessibility>1). For PT-LFC, we employed another in-house Python script to filter reads mapped to CTCF motifs and their flanking regions. Composite edit fractions were calculated for each position relative to the motif center. Using the resulting composite edit fraction curve, we defined signature peak areas (-39 to -14 and 14 to 39 positions relative to the motif center) and trough areas (-164 to -122 and 122 to 164 positions relative to the motif center). PT-LFC was then calculated as the log fold change between the edit fraction of the peak and trough regions.

**Modelling DddSs bias with deep learning**

We implemented a convolutional neural network (CNN) model with two convolutional and two fully connected layers to predict the observed DddSs editing counts based on DNA sequence. To train the model, we generated genome-wide single-nucleotide resolution signal from the E. coli data by counting the DddSs edit events. We split the E. coli genome as training (from 1 to 3641652) and validation (from 3641652 to 4641651) dataset to avoid overfitting. We binned the genome as 128 bp windows and encoded the DNA sequence with one-hot encoding. The model was trained with mean squared error (MSE) and Adam optimizer (learning_rate=0.003, weight_decay=0.0001) with 200 epochs. We reduced the learning rate when the validation error stopped decreasing for ten epochs and saved the model with the lowest validation error. We took the DNA sequence from hg38 as input and predicted the DddSs edit counts using the model described above, generating an expected ACCESS profile for human genome.

**Benchmarking dataset for transcription factor binding sites prediction**

We collected 1333 motifs derived from 409 ChIP-seq peaks in K562 from Factorbook[28]. For each of the motifs, we ran FIMO algorithm (--thresh 0.0001) to obtain its motif-based predicted binding sites (MPBSs) using the reference genome hg38. These MPBSs were used as our training data. To obtain the true labels for each motif, we intersected the MPBSs with the corresponding ChIP-seq peaks, and considered MPBSs supported by ChIP-seq peaks as true positives, and MPBS without ChIP-seq evidence as true negatives.

**Prediction of TF binding sites using ATAC-seq and ACCESS-ATAC-seq data**

To fairly compare the performance of predicting TF binding sites between ATAC-seq and ACCESS-ATAC-seq data, we sub-sampled the alignment BAM files using samtools (v1.15.1) to have the same number of reads (10M) between ACCESS-ATAC-seq (concurrent version) and ATAC-seq. Next, we used Genrich downloaded from https://github.com/jsh58/Genrich (v0.6.1) to identify peaks (-j -d 150 -D -p 0.05) after sorting the BAM files by reads name. We removed the peaks that overlapped with hg38 blacklist regions. Next, we used bedtools (v2.31.1) to identify the shared peaks between ACCESS-ATAC-seq and ATAC-seq. For each motif, we only considered the binding sites within the shared peaks. We split the chromosomes into training (chr2, chr3, chr5, chr7, chr10, chr11, chr12, chr13, chr14, chr15, chr16, chr17, chr18, chr19, chr21, chr22, chrX), validation (chr6, chr8, chr20) and test (chr1, chr4, chr9). We only kept the motifs that have at least 5% true positives in training, validation and test chromosomes to make the evaluation robust. After filtering, we obtained a total of 1076 motifs. We trained five models for each motif using different input data (DNA-only, DNA+ATAC-only, DNA+ATAC, DNA+ACCESS and DNA+ACCESS+ATAC). To account for the bias in ATAC-seq, we downloaded pre-calculated Tn5 motif bias from https://zenodo.org/records/7121027#.ZCbw4uzMI8N. For each model, we trained it using the binary cross entropy (BCE) loss by 100 epochs with a minibatch size of 48 using the Adam optimizer (learning_rate=0.003, weight_decay=0.0001). Models with the lowest validation error were saved and evaluated on the testing chromosomes.

#### *De novo* TF footprint detection

To detect TF-DNA interaction, we first smoothed the observed and predicted DddSs edit counts by averaging the single-nucleotide signal with 5bp window. Next, we computed a footprint score for each nucleotide by comparing its signal with the left and right flank regions as follows:

$$FS(i) = \frac{Signal_{Left} + Signal_{Right}}{Signal(i)}$$

A higher footprint score indicates a higher probability of the nucleotide being bound by a TF. To account for enzyme bias, we also computed the score using predicted DddSs edit counts, denoted as background footprint score. To quantify the significance, we fitted a normal distribution using the background footprint score and estimated the p-values which were further adjusted using Benjamini-Hochberg (BH) approach to account for multiple tests. We used a cutoff of 0.01 to select footprints. To compare the performance of ACCESS and ATAC, we also used Tn5 insertions to identify TF footprints as described above. For evaluation, we overlapped the detected footprints with ChIP-seq peaks from 409 TFs in K562 and computed the Jaccard Index. A higher value presents a better consistency between TF footprints and ChIP-seq peaks.

#### Pre-processing Factorbook motifs

We retrieved motif position weight matrices (PWMs) from Factorbook and used FIMO to predict motif-based predicted binding sites (MPBSs) for 717 and 1,333 motifs in the HepG2 and K562 cell lines, respectively. The motifs were filtered using a master list of DNA-binding transcription factors (TFs). For motifs with multiple associated PWMs, we selected the PWM with the lowest E-value, followed by the highest count of motif sites. For each DNA-binding TF, a motif site was designated as "bound" if it overlapped with any narrow peaks in the corresponding ChIP-seq experiment. We further refined the bound motif sites by requiring each site to have a FIMO p-value less than 0.0001, a ChIP-seq q-value (adjusted p-value) less than 0.05, and at least 50 ACCESS-ATAC-seq reads mapped to it. This filtering resulted in 119 TF motifs for the HepG2 cell line and 140 TF motifs for the K562 cell line.

#### Occupancy Pattern Inference by Editing (OccuPIE) model overview

OccuPIE is a convolutional neural network model designed to infer the occupancy state of TFs at single-allele resolution. OccuPIE employs a sequential architecture comprising three convolutional layers followed by two fully connected layers. It takes a 301 × 7 matrix as input to represent a 150-nt flanking region on either side of the motif site. The input matrix adopts a one-hot encoding format, with four channels indicating the presence of A, T, G, and C in the genomic sequence, two channels indicating C-to-T and G-to-A edits in the ACCESS-ATAC-seq read, and one channel representing the read coverage across the 301 positions. This design enables OccuPIE to leverage both the sequence and edit patterns to predict the probability of a TF-associated allele (read) being in one of three bounding states: bound, unbound-accessible, or unbound-inaccessible. Details on the model architecture and the input data structure are provided in the Supplementary Text.

**TF-specific binding state assignment**

Binding states were assigned to reads mapped to ChIP-seq-bound motifs using predefined rules that compared edit fractions in the footprint and immediate neighboring peak regions against TF-specific thresholds. Bound reads were characterized by high editing activity in the peaks and low editing in the footprint, while unbound-accessible reads displayed high editing in both the peaks and the footprint. Unbound-inaccessible reads were defined by low editing in both regions. The footprint and peak regions were determined based on the composite edit patterns specific to each TF. To evaluate the assigned states, we calculated the Spearman correlation between the fraction of bound reads and the log-transformed normalized ChIP-seq scores for the motif sites. Normalized ChIP-seq scores were computed by scaling raw signals to a 0–1 range. Thresholds for the peaks and footprints were derived from the composite motif edit profile and optimized to maximize the correlation between bound read fractions and log-transformed normalized ChIP-seq scores. Bound motif sites on Chromosome 1 were designated for defining the ranges of the footprint and peaks, as well as for optimizing the edit fraction thresholds. TF motifs in HepG2 and K562 cell lines achieving a Spearman correlation (rho) greater than 0.1 with the assigned states were selected for modeling with OccuPIE. A detailed description of the range identification and state assignment processes is provided in the supplementary text.

**OccuPIE model training and evaluation**

For each TF, bound motif sites from 18 chromosomes (chr3–chr11, chr13–chr21) were used for training, while sites from three chromosomes (chr2, chr12, chr22) were reserved for testing. Models were trained for up to 500 epochs with an adaptive learning rate and early stopping enabled. To assess the contribution of ACCESS-ATAC-seq data, a baseline model was trained using only the genomic sequence input channels. With the testing data, read states were predicted by both the full model and the baseline model, and their performances were compared based on weighted F1 scores and the area under the precision-recall curve (AUPRC). Additionally, model performance was similarly evaluated through correlation analysis with log-normalized ChIP-seq scores. Instead of using the fraction of bound reads to indicate the binding status of a TF, the mean bound probability of all reads mapped to the site was used to represent the bound probability at the motif level. Models achieving a Spearman correlation (rho) greater than 0.2 (p-value < 0.05) were considered to have good concordance with ChIP-seq evidence and were included in downstream single-allele analyses. Further details on training parameters can be found in the supplementary text.

**Co-occupancy analysis and periodicity detection**

We systematically identified pairs of adjacent bound motif sites for any two transcription factors (including self-pairs) with motif centers located within 100nt of each other. Motif pairs with overlapping footprint ranges were excluded from the analysis, as OccuPIE cannot reliably distinguish spatial differences between overlapping sites. To ensure precise center disposition calculations, motif centers were adjusted by the mean position of the footprint range relative to the FIMO-predicted center. For a given adjacent motif pair between $TF_A$ and $TF_B$ with N shared reads, we used TF-specific OccuPIE models to predict the read-level bound probability for both TFs. For the $i$-th shared read, the bound probabilities of $TF_A$ and $TF_B$ can be denoted as $P(Bound \mid A_i)$ and $P(Bound \mid B_i)$ respectively. The "observed" co-occupancy probability between the two TFs was calculated as:

$$P_{observed} = \frac{1}{N}(\sum_{i=1}^{N}(P(Bound \mid A_i) \times P(Bound \mid B_i)))$$

However, co-occupancy can also occur by random chance, particularly if the motif-level bound probability of either TFs is high. This "expected" co-occupancy probability was calculated as:

$$P_{expected} = (\frac{1}{N}\sum_{i=1}^{N} P(Bound \mid A_i)) \times (\frac{1}{N}\sum_{i=1}^{N} P(Bound \mid B_i))$$

To account for co-occupancy due to random chance, we calculated the difference between the observed and expected co-occupancy probabilities ($P_{observed} - P_{expected}$) to quantify the extent to which observed co-occupancy deviated from expected values. Motif pairs where either TF had an extreme motif-level bound probability below 0.01 or above 0.99 were excluded from further analysis. For each pair of TFs, observed – expected co-occupancy values were pooled, and a one-sample T-test was performed to assess whether the mean difference was significantly different from zero. The p-values from these tests were corrected for multiple comparisons using the Bonferroni method, with a significance threshold of alpha = 0.05.

For each TF, we applied the fast Fourier transformation (FFT) algorithm to detect periodic patterns in the observed – expected co-occupancy values between the TF and all other TFs across center displacements ranging from -100nt to 100 nt. To enable comparisons of periodicity across TFs, observed – expected values were normalized by scaling the 5th percentile to 0 and the 95th percentile to 1. Center displacement values were binned to the nearest integer, and the median observed – expected value was calculated for each bin. This pre-processing step ensured that the co-occupancy values were evenly distributed, optimizing the data for FFT analysis. FFT transformed the median observed – expected values into a spectrum of component frequencies, each characterized by an amplitude. The significance of these frequencies was evaluated using a randomization test. Null distributions for each frequency were generated by shuffling the normalized observed – expected values and performing FFT across 2000 iterations. The p-value for each frequency, representing the likelihood of observing an amplitude equal to or greater than the actual amplitude under the null hypothesis, was calculated as the proportion of randomization iterations in which the randomized amplitude exceeded or matched the real amplitude. We restricted the period detection range to 8–13nt to exclude excessively high or low frequencies. The dominant period of a TF was defined as the period corresponding to the most significant frequency within the detection range. P-values across TFs were corrected for multiple testing using the Bonferroni method (alpha = 0.05). Periodicity strength was defined as the ratio of the dominant frequency amplitude to the 95th percentile of its null distribution.

**Single-cell ACCESS-ATAC computational analysis**

We used barcoded Tn5 to assign each sequenced read to the corresponding CRISPR-KO and WT experiment. For each experiment, we grouped the cells and used the iterative approach as described above to align the reads to reference genome hg38. We filtered the BAM files by only retaining the properly paired mapped reads. Next, we converted the BAM files to fragment files and removed the duplicated reads with same start and end positions. To control the data quality, we used the package ArchR[58] to filter low-quality cells for each experiment independently based on TSS enrichment (> 2 for WT; > 2.5 for sgATXL7L3 and sgJMJD1C; > 3 for sgZBTB7A and > 4 for rest of the KO experiments) and the number of unique fragments (> 1000 and < 100000), obtaining 26147 high-quality ACCESS-ATAC single cells. We used the functions addGroupCoverages to create sample-specific Tn5 insertion coverages and addReproduciblePeakSet (cutOff = 0.01) to identify peaks for each sample. All peaks were merged to create a union peak set (n = 72166), and a count matrix was constructed with the function addPeakMatrix.

Next, we used the package Signac[59] to analyze this count matrix. Specifically, we used the functions RunTFIDF to normalize the count matrix and RunSVD to perform dimensionality reduction. For visualization, we used the function RunUMAP to generate a 2D embedding after excluding the first component which has been shown to be highly correlated with the sequencing depth of the cells. We clustered the cells using the functions FindNeighbors and FindClusters, identifying two populations (denoted as C1 and C2). We split the BAM file for each experiment based on the clusters and merged the files from all conditions to create a bulk BAM file for each cluster. To annotate these two clusters, we downloaded ATAC-seq data of HepG2 and K562 from ENCODE. We sub-sampled the BAM files to 100M reads and computed the genome-wide Spearman correlation based on Tn5 read coverage.

**Supplementary Text**

***Ddd sequences used for in vitro transcription and translation.***

T7 promoter stub

Ddd enzyme sequence

T7 terminator stub

CrDa01_T7

AGGGCTTAAGTATAAGGAGGAAAAAATATGGGTGCGGGAGCGGAAGCGATATACAAAGCTAGTAAG
GACGGGTTCAAGGAGGCGAAGAATGCGTTGCGCACTGTAAAGGACGAGAAGAACGGATTTGTAGA
CGGAGTGGCCGATACCAACGCTCCTTGGAAGGCGATAGATGACTACAGAAACCAGCGCCAGGGGT
TAGAAGTATTGCCGGAGGATTATGAGTTTATCAAAGGTGATGGAAAAAACACAGTGGCATTTAATGA
TACCTGCGACAAACGTTACTTCGGTGTCAACTCGACATTGCGCACAGATGCTGAGAAAGGTCTTGC
AAAGAAATATTTAACAAGCTGGTTGAAGAGGGTTACTTCCCGAAGGGAAGCATCTACCCTAGAGG
AAAAGCTCAATTTATCACACATGCAGAGGGTTACACGCTGATTAAGGCATACGAAGAAAATGGCAT
CGATATTGGAAAATCAGTTACGATATACGTTGATCGTCCGACTTGTGGATTTTGCCAAAATAACCTGC
CCAAGTTGCAGGCAAGCATGGGCATTGACGTGTTGACTGTCATCAATAAAAATGGCGATGTCTTCCG
GTGCGATTTGCGGAAATACACAAGTATGACGGTTTCTGACGTGAACGCAGAGAAACTGATAAAGAT
AAAATCTAAGTAGGGTTAACTAGCATAACCCCTCT

CseDa01_T7

AGGGCTTAAGTATAAGGAGGAAAAAATATGTACGATGCGTTCGGTAACGTTCGTAACCAAAAGGAA
ACCCATCACAATCGCATTTTATACACGGGTCAACAATACGACAAAGAATCAAATCAATACTATCTGC
GGGCTAGATACTACAACCCAACCCTTGGCCGTTTCACTCAAGAGGATGTTTACCGTGGGGCGGGGT
TAAATTTGTACGATTATTGCAAAGGCAACCCGGTTATATGGTACGACCCTTCAGGCTACGAAAATTGT
AAGATAGATAAAGGGAATGTATCCAATAAAGAGAACGGATTCGAGGATGAGATCATCCAGGCCAAG
ATCGCCCAACAAACCGCCTTCGAATTGGCGGAAAAGTATGGGTATAACGGAGCGCCACCGAAGAG
AAAAACAGTTGCCTCAGATGGGGAGATAAATACGCTGTCTGGGTGGAAAAAATTAAAGGATAACAC
CGATTTCATCCGTGTGTCGCCCGAGGATATTATTGACAAATCTCAGGAGATAGGACACAATCTTAGA
AACGCTGGAGCAAACGATCAGGGCATTAAGGGTAAATACAACGCTTCTCACGCTGAGAAACAATTG
TCTTTAAAGACTGACAAACCTATTGGCATCTCTCAACCAATGTGCCAGGATTGCCAGAATTACTTTC
GGTGTTTAGCTATTTCCGAAAGTAAGAACTTCGTTACTGCAGATACCAATATGGTACGTATTTTCAAG
CCGGACGGTTCCGTGATCAATTACAAGTCCGACGACTCCTTTTCGATTATAAAAATAAAGAACAAGA
TCAAATAAGGTTAACTAGCATAACCCCTCT

Ddd_Ss_T7

AGGGCTTAAGTATAAGGAGGAAAAAATATGATTAGTTTACCAGAATATGATGGAACAACTACACATG
GAGTATTAGTTTTAGATGATGGAACACAAATAGGATTTACATCAGGTAATGGAGACCCACGGTATACT
AATTATCGTAATAATGGTCATGTGGAACAGAAGTCAGCATTATATATGAGAGAAAATAATATATCCAAT
GCAACAGTATATCACAATAATACAAATGGTACTTGTGGTTATTGTAACACTATGACAGCAACTTTCTT
GCCAGAAGGAGCAACTTTAACTGTAGTTCCTCCTGAGAACGCCGTTGCAAATAATAGTAGGGCTATA
GATTATGTTAAAACGTATACGGGAACAAGTAATGACCCGAAGATAAGTCCAAGATATAAAGGAAACT
GAGGTTAACTAGCATAACCCCTCT

FlDa01_T7

AGGGCTTAAGTATAAGGAGGAAAAAATATGGTTTATGATAAACCGGAGCCTATCAAAAACCTGATCA
CATGGGTCTACGAAGGTGGCTCTTTCGTGCCGTCCGCGAAGATTATTGGCGAACATAAATTCTCTATA
ATAAATGACTACATAGGCCGTCCAATCCAAGTCTATAATGAGGTAGGGGATGTCGTGTGGGAGACCG
ACTACGATATCTATGGTGGGCTTCGTAACCTGAAAGGAGACAAGTCTTTTATACCTTTCCGGCAATTG
GGCCAATACGAGGATGTAGAAACTGGCTTATACTATAACCGCCATCGTTATTACAATCCCGAAAGCG
GTGGGTATATTAGCCAGGATCCTATAGGACTGCTTGGTGGTAGTGCGTCTTACAAGTACGTCCATGAT
TGTAACAACTGTGTAGATATTTTCGGGTTGAATCCTGTAATCTTTTCGGAAGAATTGAGTAAGATCGC
ACAGGAAGCGCATAATGTCCTGTTAGAACCCGGAAAGTCTCCTCGTGGATTTAACAACTCTACGGT
CAGTGTAGCAAAGGTCGATGTCAACGGAGTTAGCACTTTATACGCATCGGGAAATGGAGCGTCGCT
TTCTCCCGCCCAGCGTACTAAGTTGGTAGAATTAGGGGTCCCCGAAGAGAACATATTCAGTGGCAA
ACGGTTCAAAGAAATCATTGATGGGGACACAGGAACCCTTACCAAGCTGTCTAACCACGCTGAAAG
AGTGATCGAACGCAATATACCGAAGGACGCCAGCATAAAAGAGTGGGGGATTAGTTGGGCAAGTAA
GCAGAAAAATGAAATGTGCAACAACTGCAAGACCCACTTTGGATGCAAATAGGGTTAACTAGCATA
ACCCCTCT

MGYPDa01_T7

AGGGCTTAAGTATAAGGAGGAAAAAATATGTATTTCGATGGGGAGACCGGCCTGCATTACAATCGTT
TCCGTTATTACGACCCGGTGGTGGGCCGCTTTGTTCACCAAGACCCCATTGGATTTGGCGGCGGGA
ATAATTTCTACATATACGGCCCGAACAGTGGCAGTTGGTATGACCCTTTTGGCCTGGCAAAGAGACC
TCCCCATAAATTAAAAGCTAAGACTAAGGACGAGAACGGGAATGTGAAAAGTGAACGCGACTACG
TCTCTGGCGGAATGACTGAAGAGGATAAAAAACTTGGGTACCCACTTTGCTCCCTGGTTACGCATAC
GGAGAGAAAGGCCCTGAAGGAGGATGACTATTCACCGACGGACACGATTGAAATGCACGGGGAGT
ATGCTCCCTGTTCGCACTGCAAAGGTGCGATGAATACAGCGGTCGATTCTGGTAGAGTCGCACGGA
TCTTATATTACTGGAAGGGAAAGATCTGGGAGGCAGGCGCTGCTGCTCGCAAACTTAGAAATAAGA
AAAAGCGCAGCGCAGGAATGTGTTCCAATGATTAGGGTTAACTAGCATAACCCCTCT

MGYPDa029_T7

AGGGCTTAAGTATAAGGAGGAAAAAATatgGATTTCCATACGTATTATGTTGGTACAGAAAGTGTATTG
GTCCACAACAACGGGGGAAACTGTATGAAAACGGCTTCCGCGCTGGCCCAAGGTGGTTCAAAGGG
GGGATCTACCGCAATCAATCTGCCGGAATACGATGGAAAGACGACACACGGGGTCTTGGTCTTGGA
CAATGGGACTCAAGTTCAACTTGTGTCAGGCAATGGAGATCCTCGGTACACGAATTATCGCAACAAT
GGGCATGTCGAACAGAAAGCAGCGATCTATATGCGGGAAAATAACATATCAAACGCGACCGTTTAC
CATAATAACACGAATGGCACTTGCGGCTACTGTAACACCATGACCGCTACTTTCTTACCAGAGGGTG
CGACTCTGACAGTAGTTCCCCCAAAGAACGCCGTTGCGAATAACAGCCGGGCTATTGCTTACGTCA
AAACATATACCGGGACTAGCAATGACCCTAAAATGAGTTCGAGATACAAAGGCAACtagGGTTAACTA
GCATAACCCCTCT

T7 promoter extension fw primer GCGAATTAATACGACTCACTATAGGGCTTAAGTATAAGGAGGAAAA

T7 terminator extension rv primer CCAAATAAACCCCTCCGTTTAGAGAGGGGTTATGCTAGTTAACC

***DddSs and DddSs-I insert sequences used to clone pETDuet_DddSs-DddSs-I***

6x His tag

DddSs

DddSs-I

ATGGGCAGCAGCCATCACCATCATCACCACAGCCAGGATCCGATTAGTTTACCAGAATATGATGGAA
CAACTACACATGGAGTATTAGTTTTAGATGATGGAACACAAATAGGATTTACATCAGGTAATGGAGA
CCCACGGTATACTAATTATCGTAATAATGGTCATGTGGAACAGAAGTCAGCATTATATATGAGAGAAA
ATAATATATCCAATGCAACAGTATATCACAATAATACAAATGGTACTTGTGGTTATTGTAACACTATGA
CAGCAACTTTCTTGCCAGAAGGAGCAACTTTAACTGTAGTTCCTCCTGAGAACGCCGTTGCAAATA
ATAGTAGGGCTATAGATTATGTTAAAACGTATACGGGAACAAGTAATGACCCGAAGATAAGTCCAAG
ATATAAAGGAAACTGAGCGGCCGCATAATGCTTAAGTCGAACAGAAAGTAATCGTATTGTACACGGC
CGCATAATCGAAATTAATACGACTCACTATAGGGGAATTGTGAGCGGATAACAATTCCCCATCTTAGT
ATATTAGTTAAGTATAAGAAGGAGATATACATATGTTAGTAGAACATTTTATGGGACAAAAGGAATGT
GATAGTCTTGAAGAATTGAGAGAAGTTTTAAGCGAGAGGACAGAAAAAGGTGTTAATGAGTTTATT
ATATCAACGCATGAACAATTTCCGTATATGATAATGTCTGTGAAGGAAAAATATGCATGTTTAAGCTAT
TTTCGAGAAGAAGATGACCCAGGGTATTCTTCAGTTAATGCTAATCCTGTTTTGGATGCAGATGGTAT
TAGTATATTTTACACAAATACTGATAGTGAAGAAATAGAAGTTGCAAATTATTCAATTGTAAAAATTG
AAGATGCAGTGTCTGCAGTTGAGGAATTTTTTGAAACACTACAATTACCAAAATGTATTGAATGGGA
AGAATTATAG

***dsDNA deamination assay***

1. Substrate preparation:
   a. Resuspend the following oligos at 100 uM in TE buffer.

Ddd_substrate_fw_FAM /56-FAM/AAAAAAAAAAAAAAAAAAAAAAAAAAAAAAAAAAAAAAAAAAAAAAAAGATCAAAAAAAAAAAAAAAAAAAAAAAAAAAAAAAAAAAAAAAAAAAAAA

Ddd_substrate_rv TTTTTTTTTTTTTTTTTTTTTTTTTTTTTTTTTTTTTTTTTTTTTTTTGATCTTTTTTTTTTTTTTTTTTTTTTTTTTTTTTTTTTTTTTTTTTTT

   b. Mix the following (scale as necessary):

2.5 uL Ddd_substrate_fw_FAM

2.5 uL Ddd_substrate_rv

5 uL IDT Nuclease Free Duplex Buffer (cat# 11-05-01-12)

   c. Anneal oligos

In PCR block, heat to 95 deg C for 5 min, then cool to 25 deg with cooling rate of 1 deg/min (set this manually). Then cool to 4 deg C for temporary storage. Move to -20 for longer term storage. Note that this will produce 25 uM dsDNA Ddd substrate.

2. Deamination assay
   a. Make 5x Deamination buffer (100 mM Tris-HCl pH 7.4, 1M NaCl). Store at room temperature.

   b. Mix the following to create a 10 uL deamination reaction (can be made as mastermix omitting Ddd enzyme and dH2O). We recommend testing Ddd enzymes in the range of 50 nM to 5 uM to determine concentration with ~50% activity. We have used DddA at 500 nM and Ddd_Ss at 250 nM based on titration results. Include a control reaction with no Ddd enzyme.

2 uL 5x Deamination Buffer

1 uL 10 mM DTT

0.4 uL 25 uM dsDNA substrate (see above)

XX uL Ddd enzyme

XX uL dH2O (to 10 uL final volumen)

   c. Incubate reaction in PCR machine at 37C for 1 hour
   d. Store on ice and move directly to step below.

3. Detection of deamination:
   a. Perform DpnII digest as follows:

1 uL 10X DpnII Buffer (NEB)

| | |
|---|---|
| 0.5 uL | DpnII (NEB) |
| 1.5 uL | dH2O |
| 3 uL | Deamination reaction from above |

b. Incubate at 37C for 1hr (PCR machine is best)
c. Run on D1000 Tapestation. Calculate fraction deaminated as follows:

[Concentration of 90-nt product, nM]/[(Concentration of 45-nt product, nM/2) + (Concentration of 90-nt product, nM)

d. We have found that the concentration of Ddd enzyme producing ~50% deamination in this dsDNA deamination assay is optimal as final Ddd enzyme concentration for ACCESS.

**Identifying motif features**

The composite edit fraction for bound motifs is smoothed using a 15nt rolling average to reduce noise and highlight meaningful patterns. Here we denote the relative position to motif center as $x$, the edit fraction and the smoothed edit fraction at position x as $Ef(x)$ and $Ef_{smooth}(x)$ respectively. We calculated two key edit fraction values:

- minimum footprint edit fraction: $Ef_{\text{f\_min}} = \min\big(Ef(x)\big), x \in [-12, 12]$
- minimum of the left peak and right peak maximum edit fractions : $Ef_{\text{p\_max\_}min} = \min \begin{cases} \max\big(Ef_{smooth}(x)\big), x \in [-75, -14] \\ \max\big(Ef_{smooth}(x)\big), x \in [14, 75] \end{cases}$

Next, we calculate the feature edit fraction ratio $r(x)$ as:

$$r(x) = (Ef(x) - Ef_{f\_min})/(Ef(x) - Ef_{p\_\text{max}\ \_min})$$

The footprint positions $P_f$, the left peak positions $P_{lp}$ and the right peak positions $P_{rp}$ are then identified as:

- $P_f$: any position where $r(x) < 0.55$ for $x \in [-12,12]$
- $P_{lp}$: any position where $r(x) > 0.67$ for $x \in [-100, \min(P_f)]$
- $P_{rp}$: any position where $r(x) > 0.67$ for $x \in [\max\big(P_f\big), 100]$

**OccuPIE training data states assignment**

For a given ACCESS-ATAC-seq read mapped to a motif site, we calculate the mean edit fraction of the footprint positions as $\mu_{footprint}$ , the mean edit fraction of both the left and right peak positions as $\mu_{peak}$, as well as the mean edit fraction of all positions (within ±150nt from motif center) as $\mu_{global}$. Then the read's state can be assigned given the corresponding thresholds ($Thres$) for each value:

- Unbound-inaccessible, if $\mu_{peak} < Thres_{peak}$ and $\mu_{global} < Thres_{global}$
- Bound, if read is not unbound-inaccessible and $\mu_{footprint} < Thres_{footprint}$
- Unbound-accessible, if read is neither unbound-inaccessible nor bound

The peak and footprint thresholds are further determined by the corresponding weights ($w$):

$$Thres_{peak} = \mu_{peak} \cdot w_{peak} + \mu_{footprint} \cdot (1 - w_{peak})$$
$$Thres_{footprint} = \mu_{footprint} \cdot w_{footprint} + \mu_{peak} \cdot (1 - w_{footprint})$$

The global threshold is calculated using the mean edit fraction and the edit fraction standard deviation of unbound motifs (within ±150nt from motif center):

$$Thres_{global} = \mu_{unbound} + 2\sigma_{unbound}$$

Finally, the weights for peak and footprint are systematically determined for each different motif. For a given motif, we iteratively try $w_{peak}$ and $w_{footprint}$ values from -2 to 2 with a step of 0.1. In each iteration, we assign states to reads and calculate the sparman correlation between the resulting bound read fractions and the log transformed normalized ChIP-seq scores. The weight combination that achieve the highest correlation is used to assign states for the training data.

**OccuPIE input matrix data structure**

Each read mapped to a motif site, representing a single allele, is processed into a 301 × 7 multi-channel one-hot encoded input matrix. This matrix is centered on the motif site, extending 150 nucleotides upstream and downstream. The input matrix consists of the following 7 channels:

- Base Channels (4): These channels encode the A, T, G, and C nucleotides in the reference sequence.
- Edit Channels (2): These channels encode "C-to-T" and "G-to-A" edits observed in the ACCESS-ATAC-seq read.
- Coverage Channel (1): This channel encodes the read coverage relative to the motif center.

**OccuPIE model architecture and training parameters**

The OccuPIE model is built on the tensorflow frameowork with a sequential architecture. The model is mainly comprised of three convolutional layers and two dense layers. The convolutional layers contain 16, 32 and 64 1d filters with a fixed filter size of 12.

***ACCESS-WGS protocol***

1. Prepare stock solutions and buffers.

The following buffers can be made in advance and stored.

**RSB:** 10 mM Tris-HCl (pH 7.5, Invitrogen, cat. no. 15567027), 10 mM NaCl (Invitrogen, cat. no. AM9759) and 3 mM $MgCl_2$ (Invitrogen, cat. no. AM9530G) in nuclease-free dH2O. RSB can be made in bulk and stored at 4 °C long-term.

**10% IGEPAL** in nuclease-free dH2O (Sigma, cat. no. I3021). Mix well and store at 4 °C long-term.

**1% Digitonin** in nuclease-free dH2O (Invitrogen, cat. no. BN2006). Mix well and store at 4 °C long-term.

**10% Tween-20** in nuclease-free dH2O (Bio-Rad, cat. no. 1610781). Mix well and store at 4 °C long-term.

**2x ATAC-seq buffer**: 20 mM Tris HCl (pH 7.5), 10 mM MgCl2, 20% Dimethyl Formamide.

The following buffers should be made on the day of the experiment:

**ATAC lysis buffer**: Add 0.1% IGEPAL (Sigma, cat. no. I3021, 1:100 from 10% IGEPAL stock solution), 0.01% digitonin (Invitrogen, cat. no. BN2006, 1:100 from 1% digitonin stock solution), and 0.1% Tween-20 (Bio-Rad, cat. no. 1610781, 1:100 from 10% Tween-20 stock solution) to RSB. Make 150 uL per sample to be tested. Detergent percentages reported are final concentrations.

**RSB + 0.1% Tween-20:** Add 0.1% Tween-20 (Bio-Rad, cat. no. 1610781, 1:100 from 10% Tween-20 stock solution) to RSB. Make 1.2-2 mL per sample to be tested. Detergent percentages reported are final concentrations.

2. Prepare intact nuclei.

- We have performed ACCESS-ATAC using 2.5*10^5 cells in 25 uL reaction volume or 5*10^5 cells in 50 uL reaction volume. It is likely that the protocol would be successful at a range of cell concentrations.
- For non-adherent cells, spin and resuspend in 1 mL ice-cold PBS. Pipet up and down to mix well. Perform cell counting while keeping cells on ice.
- For adherent cells, trypsinize, quench, spin, and resuspend in 1 mL ice-cold PBS. Pipet up and down to mix well. Perform cell counting while keeping cells on ice.
- Using the live cell count, pipet desired cells into one centrifuge tube. Add PBS up to at least 1mL (if there is >100 uL based on what you transfer, that's fine).
- Spin down at 500xg for 5 min at 4 deg.
- Remove supernatant very carefully. By pipetting, thoroughly resuspend each cell pellet in 125 uL ice-cold ATAC Lysis Buffer.
- After resuspending cell pellets in the lysis buffer, incubate on ice for 3 min, and then stop lysis by adding 1.3 ml RSB + 0.1% Tween-20 to each tube.
- Centrifuge nuclei at 500 r.c.f for 10 min at 4 °C. Remove supernatant and make sure no more than 20 uL are left over. Do not disturb pellet.
- Prepare thermal block at 37 deg

3. Treat nuclei with Ddd enzyme.

   a. Prepare ACCESS treatment mix (can be made as mastermix for multiple samples). Shown for 25 uL reaction, can be scaled to 50 uL reaction volume:

1.5 uL 2x ATAC-seq buffer
1.5 uL 1% digitonin stock solution for 0.1% final concentration
1.5 uL 10% Tween-20 stock solution for 1% final concentration

1 uL UGI (NEB M0281S/M0281L)

0.5 uL Ddd enzyme titrated to provide ~50% deamination in dsDNA deamination assay

b. Gently resuspend nuclei pellet with 19 uL ACCESS treatment mix. Use pipet to measure reaction volume. If <25 uL, add RSB + 0.1% Tween-20 to final volume of 25 uL.
c. Incubate at 37 deg C for 30 min, flicking gently to mix every 10-15 minutes.
d. After incubation, immediately clean up treated DNA using Zymo Genomic DNA clean and concentrator 25 kit (D4064/D4065).

All centrifugation steps should be performed between 10,000 - 16,000 x g. Pre-heat 50 uL of Zymo Elution Buffer per sample to 50-60 deg C.
-In a 1.5 ml microcentrifuge tube, add 2 volumes of ChIP DNA Binding Buffer to each volume of DNA sample. In this case, add 50 uL DNA binding buffer. Mix briefly by vortexing.
-Transfer mixture to a provided Zymo-Spin™ Column in a Collection Tube.
-Centrifuge for 30 seconds. Discard the flow-through.
-Add 400 µl DNA Wash Buffer to the column. Centrifuge for 30 seconds. Repeat this wash step.
-Add 50 uL heated Zymo Elution Buffer (50-60 deg C) directly to the column matrix and incubate at room temperature for one minute. Transfer the column to a 1.5 ml microcentrifuge tube and centrifuge for 30 seconds to elute the DNA.

4. Tagment genomic DNA

- Dilute 100 ng of ACCESS-treated genomic DNA to a total volume of 20 uL (5 ng/uL).
- Add 20 µl of ACCESS-treated genomic DNA at 5 ng/µl (100 ng total) to a PCR tube.
- Add 25 µl of 2x ATAC-seq Buffer.
- Add 5 µl of mix of 0.5 uL Tagmentase (Tn5 transposase) - loaded (Diagenode C01070012-30/ C01070012-200) + 4.5 uL dH2O (0.5 uL loaded Tn5 + 4.5 uL dH2O)
- Pipette up and down 10 times to mix.
- Perform incubation in a PCR machine as follows: 55C for 5 minutes, hold at 10C
- After incubation, clean up treated DNA using Zymo DNA clean and concentrator 5 kit (D4003/D4004) using protocol below.

All centrifugation steps should be performed between 10,000 - 16,000 x g. Pre-heat 50 uL of Zymo Elution Buffer per sample to 50-60 deg C.

-In a 1.5 ml microcentrifuge tube, add 5 volumes of DNA Binding Buffer to each volume of DNA sample. In this case, add 125 uL DNA binding buffer. Mix briefly by vortexing.
-Transfer mixture to a provided Zymo-Spin™ Column in a Collection Tube.
-Centrifuge for 30 seconds. Discard the flow-through.
-Add 200 µl DNA Wash Buffer to the column. Centrifuge for 30 seconds. Repeat this wash step.
-Add 50 uL heated Zymo Elution Buffer (50-60 deg C) directly to the column matrix and incubate at room temperature for one minute. Transfer the column to a 1.5 ml microcentrifuge tube and centrifuge for 30 seconds to elute the DNA.

-Eluted DNA can be stored on ice temporarily or at -20 for 24-72 hours prior to library preparation.

5. Prepare ACCESS-WGS libraries for NGS.

a. Perform qPCR to determine appropriate cycle count for library prep. Per reaction volumes are:

0.375 uL 20 uM Nextera N7 primer (see below for sequences)

0.375 uL 20 uM Nextera N5 primer (see below for sequences)

7.5 uL 2x NEBNext **Q5U** Master Mix (NEB M0597L)

0.75 uL 20x EvaGreen Dye (Biotium #31000)

5.5 uL dH2O

0.5 uL purified ACCESS-treated DNA

qPCR settings:

72°C for 5min

98°C for 30 seconds

25 cycles of: 98°C for 10 seconds, 63°C for 30 seconds, 72°C for 30 seconds.

Hold at 10°C

Determine appropriate cycles using qPCR. Typically, perform cycles equivalent to qPCR Ct or add 1-2 cycles.

b. PCR Amplification of ACCESS-treated DNA. Per-reaction volumes are:

1.25 uL 20 uM Nextera N7 primer (see below for sequences)

1.25 uL 20 uM Nextera N5 primer (see below for sequences)

25 uL 2x NEBNext **Q5U** Master Mix (NEB M0597L)

22.5 uL purified ACCESS- treated DNA

Thermal Cycler settings:

72°C for 5min

98°C for 30 seconds

XX cycles (based on qPCR) of: 98°C for 10 seconds, 63°C for 30 seconds, 72°C for 30 seconds.

Hold at 10°C

c. SPRI clean up using a 1-sided procedure.

i. To the PCR reaction, add 55ul of mixed, ROOM TEMP, SPRI beads to each sample (this is a 1.1X SPRI).
ii. Vortex briefly and incubate for 5min at room temp
iii. Apply magnet to collect beads
iv. Once solution is clear, use pipette to remove supernatant
v. While still on magnet, add 180ul of 80% ETOH to each sample without mixing
vi. Incubate for 30 sec at room temperature
vii. Remove supernatant with pipette
viii. Repeat steps EtOH wash steps for a second ethanol wash
ix. Allow tubes to sit at room temp so the residual ethanol can evaporate, beads will turn from shiny to matte when dry (2-5 min), then proceed
x. With tubes off the magnet, add 20ul Elution Buffer (can be from Zymo or other company)
xi. Cap tubes and mix by vortexing
xii. Incubate samples for 5 min at room temp
xiii. Apply magnet to samples
xiv. Transfer supernatant to a fresh labeled tube.
xv. Samples are now ready for Tapestation D1000 and NGS

Sequences used in PCR amplification

| | |
|---|---|
| **Nextera N7 primers** | |
| Nextera_N701 | CAAGCAGAAGACGGCATACGAGAT TCGCCTTA GTCTCGTGGGCTCGG |
| Nextera_N702 | CAAGCAGAAGACGGCATACGAGAT CTAGTACG GTCTCGTGGGCTCGG |
| Nextera_N703 | CAAGCAGAAGACGGCATACGAGAT TTCTGCCT GTCTCGTGGGCTCGG |
| Nextera_N704 | CAAGCAGAAGACGGCATACGAGAT GCTCAGGA GTCTCGTGGGCTCGG |
| Nextera_N705 | CAAGCAGAAGACGGCATACGAGAT AGGAGTCC GTCTCGTGGGCTCGG |
| Nextera_N706 | CAAGCAGAAGACGGCATACGAGAT CATGCCTA GTCTCGTGGGCTCGG |
| Nextera_N707 | CAAGCAGAAGACGGCATACGAGAT GTAGAGAG GTCTCGTGGGCTCGG |
| Nextera_N708 | CAAGCAGAAGACGGCATACGAGAT CCTCTCTG GTCTCGTGGGCTCGG |
| **Nextera N5 primers** | |
| Nextera_N501 | AATGATACGGCGACCACCGAGATCTACAC TAGATCGC TCGTCGGCAGCGTC |
| Nextera_N502 | AATGATACGGCGACCACCGAGATCTACAC CTCTCTAT TCGTCGGCAGCGTC |
| Nextera_N503 | AATGATACGGCGACCACCGAGATCTACAC TATCCTCT TCGTCGGCAGCGTC |
| Nextera_N504 | AATGATACGGCGACCACCGAGATCTACAC AGAGTAGA TCGTCGGCAGCGTC |
| Nextera_N505 | AATGATACGGCGACCACCGAGATCTACAC GTAAGGAG TCGTCGGCAGCGTC |
| Nextera_N506 | AATGATACGGCGACCACCGAGATCTACAC ACTGCATA TCGTCGGCAGCGTC |
| Nextera_N507 | AATGATACGGCGACCACCGAGATCTACAC AAGGAGTA TCGTCGGCAGCGTC |
| Nextera_N508 | AATGATACGGCGACCACCGAGATCTACAC CTAAGCCT TCGTCGGCAGCGTC |

***ACCESS-ATAC protocol***

1. Prepare stock solutions and buffers.

The following buffers can be made in advance and stored.

**RSB:** 10 mM Tris-HCl (pH 7.5, Invitrogen, cat. no. 15567027), 10 mM NaCl (Invitrogen, cat. no. AM9759) and 3 mM $MgCl_2$ (Invitrogen, cat. no. AM9530G) in nuclease-free dH2O. RSB can be made in bulk and stored at 4 °C long-term.

**10% IGEPAL** in nuclease-free dH2O (Sigma, cat. no. I3021). Mix well and store at 4 °C long-term.

**1% Digitonin** in nuclease-free dH2O (Invitrogen, cat. no. BN2006). Mix well and store at 4 °C long-term.

**10% Tween-20** in nuclease-free dH2O (Bio-Rad, cat. no. 1610781). Mix well and store at 4 °C long-term.

**2x ATAC-seq buffer**: 20 mM Tris HCl (pH 7.5), 10 mM MgCl2, 20% Dimethyl Formamide.

The following buffers should be made on the day of the experiment:

**ATAC lysis buffer**: Add 0.1% IGEPAL (Sigma, cat. no. I3021, 1:100 from 10% IGEPAL stock solution), 0.01% digitonin (Invitrogen, cat. no. BN2006, 1:100 from 1% digitonin stock solution), and 0.1% Tween-20 (Bio-Rad, cat. no. 1610781, 1:100 from 10% Tween-20 stock solution) to RSB. Make 150 uL per sample to be tested. Detergent percentages reported are final concentrations.

**RSB + 0.1% Tween-20:** Add 0.1% Tween-20 (Bio-Rad, cat. no. 1610781, 1:100 from 10% Tween-20 stock solution) to RSB. Make 1.2-2 mL per sample to be tested. Detergent percentages reported are final concentrations.

2. Prepare intact nuclei.

- We have performed ACCESS-ATAC using 2.5*10^5 cells in 25 uL reaction volume or 5*10^5 cells in 50 uL reaction volume. It is likely that the protocol would be successful at a range of cell concentrations.
- For non-adherent cells, spin and resuspend in 1 mL ice-cold PBS. Pipet up and down to mix well. Perform cell counting while keeping cells on ice.
- For adherent cells, trypsinize, quench, spin, and resuspend in 1 mL ice-cold PBS. Pipet up and down to mix well. Perform cell counting while keeping cells on ice.
- Using the live cell count, pipet desired cells into one centrifuge tube. Add PBS up to at least 1mL (if there is >100 uL based on what you transfer, that's fine).
- Spin down at 500xg for 5 min at 4 deg.
- Remove supernatant very carefully. By pipetting, thoroughly resuspend each cell pellet in 125 uL ice-cold ATAC Lysis Buffer.
- After resuspending cell pellets in the lysis buffer, incubate on ice for 3 min, and then stop lysis by adding 1.3 ml RSB + 0.1% Tween-20 to each tube.
- Centrifuge nuclei at 500 r.c.f for 10 min at 4 °C. Remove supernatant and make sure no more than 20 uL are left over. Do not disturb pellet.
- Prepare thermal block at 37 deg

3. Treat nuclei with Ddd_Ss and loaded Tn5.

Concurrent protocol (recommended):

a. Prepare ACCESS-ATAC treatment mix (can be made as mastermix for multiple samples). Shown for 25 uL reaction, can be scaled to 50 uL reaction volume:

| | |
|---|---|
| 1.5 uL | 2x ATAC-seq buffer |
| 1.5 uL | 1% digitonin stock solution for 0.1% final concentration |
| 1.5 uL | 10% Tween-20 stock solution for 1% final concentration |

1 uL UGI (NEB M0281S/M0281L)

1.25 uL Tagmentase (Tn5 transposase) - loaded (Diagenode C01070012-30/ C01070012-200)

0.75 uL Ddd_Ss (8.33 uM stock for 250 nM final concentration)

b. Gently resuspend nuclei pellet with 20.5 uL ACCESS-ATAC treatment mix. Use pipet to measure reaction volume. If <25 uL, add RSB + 0.1% Tween-20 to final volume of 25 uL.
c. Incubate at 37 deg C for 30 min, flicking gently to mix every 10-15 minutes.
d. After incubation, immediately clean up treated DNA using Zymo DNA clean and concentrator 5 kit (D4003/D4004) using protocol below.

All centrifugation steps should be performed between 10,000 - 16,000 x g. Pre-heat 50 uL of Zymo Elution Buffer per sample to 50-60 deg C.
1. In a 1.5 ml microcentrifuge tube, add 5 volumes of DNA Binding Buffer to each volume of DNA sample. In this case, add 125 uL DNA binding buffer. Mix briefly by vortexing.
2. Transfer mixture to a provided Zymo-Spin™ Column in a Collection Tube.
3. Centrifuge for 30 seconds. Discard the flow-through.
4. Add 200 µl DNA Wash Buffer to the column. Centrifuge for 30 seconds. Repeat this wash step.
5. Add 50 uL heated Zymo Elution Buffer (50-60 deg C) directly to the column matrix and incubate at room temperature for one minute. Transfer the column to a 1.5 ml microcentrifuge tube and centrifuge for 30 seconds to elute the DNA.

Sequential protocol (not recommended):

a. Prepare ACCESS treatment mix (can be made as mastermix for multiple samples). Shown for 25 uL reaction, can be scaled to 50 uL reaction volume:

12.5 uL 2x ATAC-seq buffer

2.5 uL 1% digitonin stock solution for 0.1% final concentration

1.5 uL 10% Tween-20 stock solution for 1% final concentration

1 uL UGI (NEB M0281S/M0281L)

0.5 uL Ddd_Ss (11.63 uM stock for 250 nM final concentration in 23.25 uL final volume)

b. Gently resuspend nuclei pellet with 19 uL ACCESS treatment mix. Use pipet to measure reaction volume. If <23.25 uL, add RSB + 0.1% Tween-20 to final volume of 23.25 uL.
c. Incubate at 37 deg C for 15-30 min (see manuscript for effect of treatment time), flicking gently to mix every 10-15 minutes.
d. Add 5-10X molar excess of Ddd_Ss-I in 0.5 uL. Pipet up and down gently to mix. For example, add 0.5 uL of 59.4 uM Ddd_Ss-I for 1.25 uM final concentration in 23.75 uL final volume.
e. Incubate for 2 minutes at 37 deg C.
f. Add 1.25 uL Tagmentase (Tn5 transposase) - loaded (Diagenode C01070012-30/ C01070012-200).
g. Incubate at 37 deg C for 30 min, flicking gently to mix every 10-15 minutes.
h. After incubation, immediately clean up treated DNA using Zymo DNA clean and concentrator 5 kit (D4003/D4004) using protocol below.

All centrifugation steps should be performed between 10,000 - 16,000 x g. Pre-heat 50 uL of Zymo Elution Buffer per sample to 50-60 deg C.

-In a 1.5 ml microcentrifuge tube, add 5 volumes of DNA Binding Buffer to each volume of DNA sample. In this case, add 125 uL DNA binding buffer. Mix briefly by vortexing.
-Transfer mixture to a provided Zymo-Spin™ Column in a Collection Tube.
-Centrifuge for 30 seconds. Discard the flow-through.

-Add 200 µl DNA Wash Buffer to the column. Centrifuge for 30 seconds. Repeat this wash step. -Add 50 uL heated Zymo Elution Buffer (50-60 deg C) directly to the column matrix and incubate at room temperature for one minute. Transfer the column to a 1.5 ml microcentrifuge tube and centrifuge for 30 seconds to elute the DNA.

-Eluted DNA can be stored on ice temporarily or at -20 for 24-72 hours prior to library preparation.

4. Prepare ACCESS-ATAC libraries for NGS.

a. Perform qPCR to determine appropriate cycle count for library prep. Per reaction volumes are:

0.375 uL 20 uM Nextera N7 primer (see below for sequences)

0.375 uL 20 uM Nextera N5 primer (see below for sequences)

7.5 uL 2x NEBNext **Q5U** Master Mix (NEB M0597L)

0.75 uL 20x EvaGreen Dye (Biotium #31000)

5.5 uL dH2O

0.5 uL purified ACCESS-ATAC-treated DNA

qPCR settings:

72°C for 5min

98°C for 30 seconds

25 cycles of: 98°C for 10 seconds, 63°C for 30 seconds, 72°C for 30 seconds.

Hold at 10°C

Determine appropriate cycles using qPCR. Typically, perform cycles equivalent to qPCR Ct or add 1-2 cycles.

b. PCR Amplification of ACCESS-ATAC-treated DNA. Per-reaction volumes are:

1.25 uL 20 uM Nextera N7 primer (see below for sequences)

1.25 uL 20 uM Nextera N5 primer (see below for sequences)

25 uL 2x NEBNext **Q5U** Master Mix (NEB M0597L)

22.5 uL purified ACCESS-ATAC-treated DNA

Thermal Cycler settings:

72°C for 5min

98°C for 30 seconds

XX cycles (based on qPCR) of: 98°C for 10 seconds, 63°C for 30 seconds, 72°C for 30 seconds.

Hold at 10°C

c. SPRI clean up using a 1-sided procedure.

i. To the PCR reaction, add 55ul of mixed, ROOM TEMP, SPRI beads to each sample (this is a 1.1X SPRI).

ii. Vortex briefly and incubate for 5min at room temp

iii. Apply magnet to collect beads
iv. Once solution is clear, use pipette to remove supernatant
v. While still on magnet, add 180ul of 80% ETOH to each sample without mixing
vi. Incubate for 30 sec at room temperature
vii. Remove supernatant with pipette
viii. Repeat steps EtOH wash steps for a second ethanol wash
ix. Allow tubes to sit at room temp so the residual ethanol can evaporate, beads will turn from shiny to matte when dry (2-5 min), then proceed
x. With tubes off the magnet, add 20ul Elution Buffer (can be from Zymo or other company)
xi. Cap tubes and mix by vortexing
xii. Incubate samples for 5 min at room temp
xiii. Apply magnet to samples
xiv. Transfer supernatant to a fresh labeled tube.
xv. Samples are now ready for Tapestation D1000 and NGS

Sequences used in PCR amplification

| **Nextera N7 primers** | |
|---|---|
| Nextera_N701 | CAAGCAGAAGACGGCATACGAGAT TCGCCTTA GTCTCGTGGGCTCGG |
| Nextera_N702 | CAAGCAGAAGACGGCATACGAGAT CTAGTACG GTCTCGTGGGCTCGG |
| Nextera_N703 | CAAGCAGAAGACGGCATACGAGAT TTCTGCCT GTCTCGTGGGCTCGG |
| Nextera_N704 | CAAGCAGAAGACGGCATACGAGAT GCTCAGGA GTCTCGTGGGCTCGG |
| Nextera_N705 | CAAGCAGAAGACGGCATACGAGAT AGGAGTCC GTCTCGTGGGCTCGG |
| Nextera_N706 | CAAGCAGAAGACGGCATACGAGAT CATGCCTA GTCTCGTGGGCTCGG |
| Nextera_N707 | CAAGCAGAAGACGGCATACGAGAT GTAGAGAG GTCTCGTGGGCTCGG |
| Nextera_N708 | CAAGCAGAAGACGGCATACGAGAT CCTCTCTG GTCTCGTGGGCTCGG |
| **Nextera N5 primers** | |
| Nextera_N501 | AATGATACGGCGACCACCGAGATCTACAC TAGATCGC TCGTCGGCAGCGTC |
| Nextera_N502 | AATGATACGGCGACCACCGAGATCTACAC CTCTCTAT TCGTCGGCAGCGTC |
| Nextera_N503 | AATGATACGGCGACCACCGAGATCTACAC TATCCTCT TCGTCGGCAGCGTC |

| | |
|---|---|
| Nextera_N504 | AATGATACGGCGACCACCGAGATCTACAC AGAGTAGA TCGTCGGCAGCGTC |
| Nextera_N505 | AATGATACGGCGACCACCGAGATCTACAC GTAAGGAG TCGTCGGCAGCGTC |
| Nextera_N506 | AATGATACGGCGACCACCGAGATCTACAC ACTGCATA TCGTCGGCAGCGTC |
| Nextera_N507 | AATGATACGGCGACCACCGAGATCTACAC AAGGAGTA TCGTCGGCAGCGTC |
| Nextera_N508 | AATGATACGGCGACCACCGAGATCTACAC CTAAGCCT TCGTCGGCAGCGTC |

*OccuPIE implementation*

**Identifying motif features**

The composite edit fraction for bound motifs is smoothed using a 15nt rolling average to reduce noise and highlight meaningful patterns. Here we denote the relative position to motif center as $x$, the edit fraction and the smoothed edit fraction at position x as $Ef(x)$ and $Ef_{smooth}(x)$ respectively. We calculated two key edit fraction values:

- minimum footprint edit fraction: $Ef_{\text{f\_min}} = \min(Ef(x)), x \in [-12, 12]$
- minimum of the left peak and right peak maximum edit fractions : $Ef_{p\_\text{max}\_min} = \min\begin{cases}\max(Ef_{smooth}(x)), x \in [-75, -14] \\ \max(Ef_{smooth}(x)), x \in [14, 75]\end{cases}$

Next, we calculate the feature edit fraction ratio $r(x)$ as:

$$r(x) = (Ef(x) - Ef_{f\_min})/(Ef(x) - Ef_{p\_\text{max}\ \_min})$$

The footprint positions $P_f$, the left peak positions $P_{lp}$ and the right peak positions $P_{rp}$ are then identified as:

- $P_f$: any position where $r(x) < 0.55$ for $x \in [-12,12]$
- $P_{lp}$: any position where $r(x) > 0.67$ for $x \in [-100, \min(P_f)]$
- $P_{rp}$: any position where $r(x) > 0.67$ for $x \in [\max(P_f), 100]$

**OccuPIE training data states assignment**

For a given ACCESS-ATAC-seq read mapped to a motif site, we calculate the mean edit fraction of the footprint positions as $\mu_{footprint}$ , the mean edit fraction of both the left and right peak positions as $\mu_{peak}$, as well as the mean edit fraction of all positions (within ±150nt from motif center) as $\mu_{global}$. Then the read's state can be assigned given the corresponding thresholds ($Thres$) for each value:

- Unbound-inaccessible, if $\mu_{peak} < Thres_{peak}$ and $\mu_{global} < Thres_{global}$
- Bound, if read is not unbound-inaccessible and $\mu_{footprint} < Thres_{footprint}$
- Unbound-accessible, if read is neither unbound-inaccessible nor bound

The peak and footprint thresholds are further determined by the corresponding weights ($w$):

$$Thres_{peak} = \mu_{peak} \cdot w_{peak} + \mu_{footprint} \cdot (1 - w_{peak})$$
$$Thres_{footprint} = \mu_{footprint} \cdot w_{footprint} + \mu_{peak} \cdot (1 - w_{footprint})$$

The global threshold is calculated using the mean edit fraction and the edit fraction standard deviation of unbound motifs (within ±150nt from motif center):

$$Thres_{global} = \mu_{unbound} + 2\sigma_{unbound}$$

Finally, the weights for peak and footprint are systematically determined for each different motif. For a given motif, we iteratively try $w_{peak}$ and $w_{footprint}$ values from -2 to 2 with a step of 0.1. In each iteration, we assign states to reads and calculate the sparman correlation between the resulting bound read fractions and the log transformed normalized ChIP-seq scores. The weight combination that achieve the highest correlation is used to assign states for the training data.

**OccuPIE input matrix data structure**

Each read mapped to a motif site, representing a single allele, is processed into a 301 × 7 multi-channel one-hot encoded input matrix. This matrix is centered on the motif site, extending 150 nucleotides upstream and downstream. The input matrix consists of the following 7 channels:

- Base Channels (4): These channels encode the A, T, G, and C nucleotides in the reference sequence.
- Edit Channels (2): These channels encode “C-to-T” and “G-to-A” edits observed in the ACCESS-ATAC-seq read.
- Coverage Channel (1): This channel encodes the read coverage relative to the motif center.

**OccuPIE model architecture and training parameters**

The OccuPIE model is built on the TensorFlow framework with a sequential architecture. The input layer accepts data with dimensions corresponding to the preprocessed one-hot encoded dataset. The model includes three convolutional layers, each using ReLU activation and L2 regularization (coefficient 0.01). Filters in the first layer span the input height with a width of 15, while subsequent layers use a height of 1. The number of filters starts at 16 and increases linearly with the layer index. Each convolutional layer is followed by max pooling (1 × 2) and dropout (rate 0.25) to reduce overfitting. The convolutional layers are followed by a flattening layer, which converts the output into a one-dimensional vector for the dense layers. Two dense layers progressively reduce the node count, starting with 32 nodes and halving with each layer. Both dense layers use ReLU activation, L2 regularization, and dropout (rate 0.25). The final output layer includes three nodes (one for each class), with softmax activation to output class probabilities. The model is compiled with the Adam optimizer (learning rate 0.0001), categorical crossentropy loss, and accuracy as the primary metric.

The model was trained using a batch size of 4096 for a maximum of 500 epochs. Early stopping was employed to monitor validation loss and halt training after 12 epochs of no improvement, restoring the best weights. A learning rate scheduler reduced the learning rate by a factor of 0.5 when validation loss plateaued for three epochs, with a minimum learning rate of $1 \times 10^{-6}$. Class weights were calculated to address class imbalance, ensuring proportional weighting based on the inverse frequency of each class in the training dataset. Input data was expanded by an additional dimension to ensure compatibility with the convolutional layers.

**Supplementary Figures**

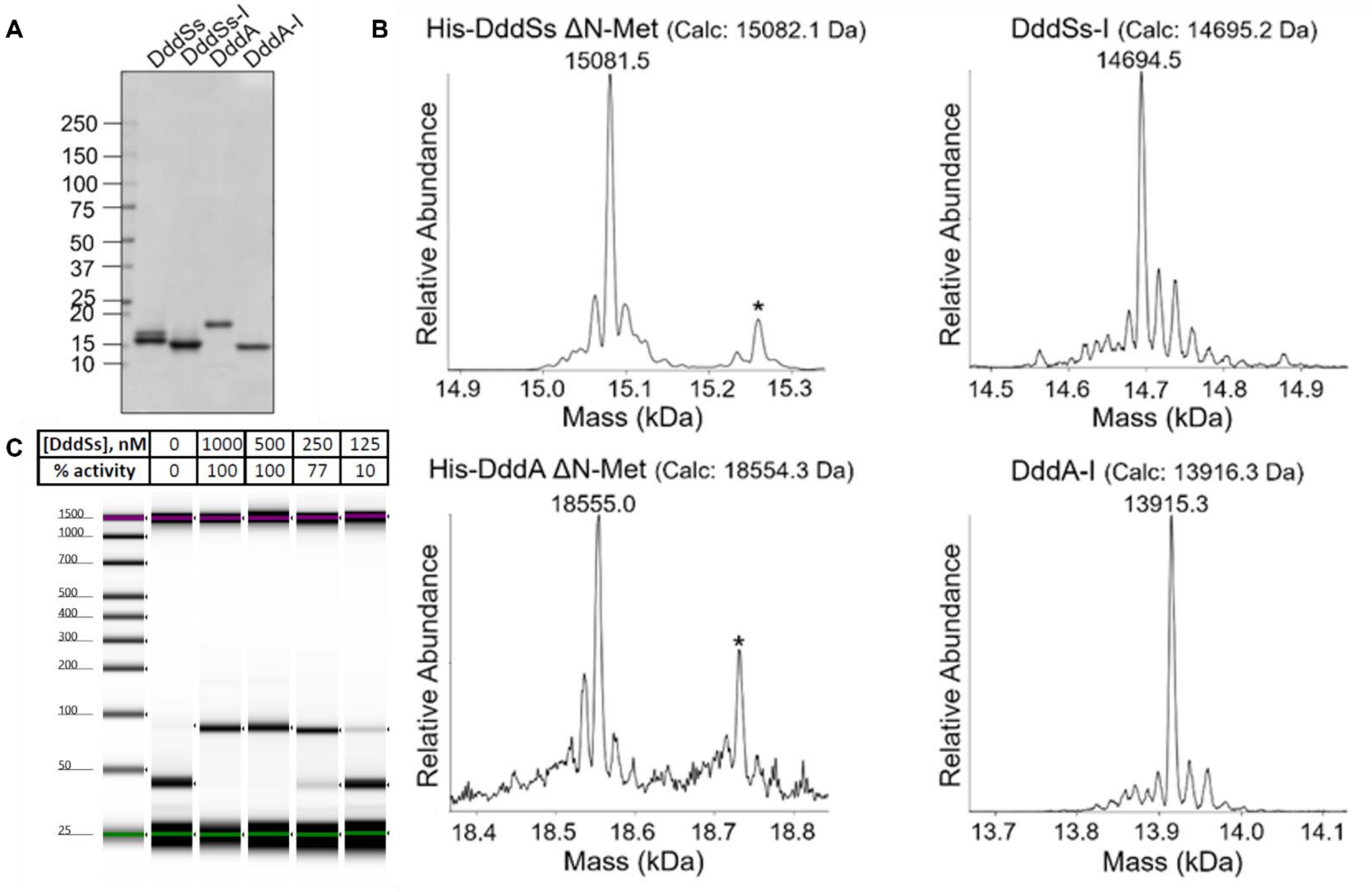


**Supplementary Figure 1: Characterization of deaminases and deaminase inhibitors by SDS-PAGE gel, ESI-MS, and deamination activity.**

(A) Coomassie-stained 4-20% SDS PAGE gel. (B) Deconvoluted mass spectra of intact protein LC-MS obtained using UniDec software. The observed masses of the products are the deconvoluted average neutral mass for each protein. The peak marked with asterisks (*) likely correspond to an adduct observed in His-tag protein purifications. (C) Agilent Tapestation gel showing results of titration of DddSs activity using a double-stranded DNA (dsDNA) deamination assay. Appearance of a 90-nt band denotes dsDNA deamination, which impedes DpnII restriction digest. DddSs concentration and deamination activity are reported.

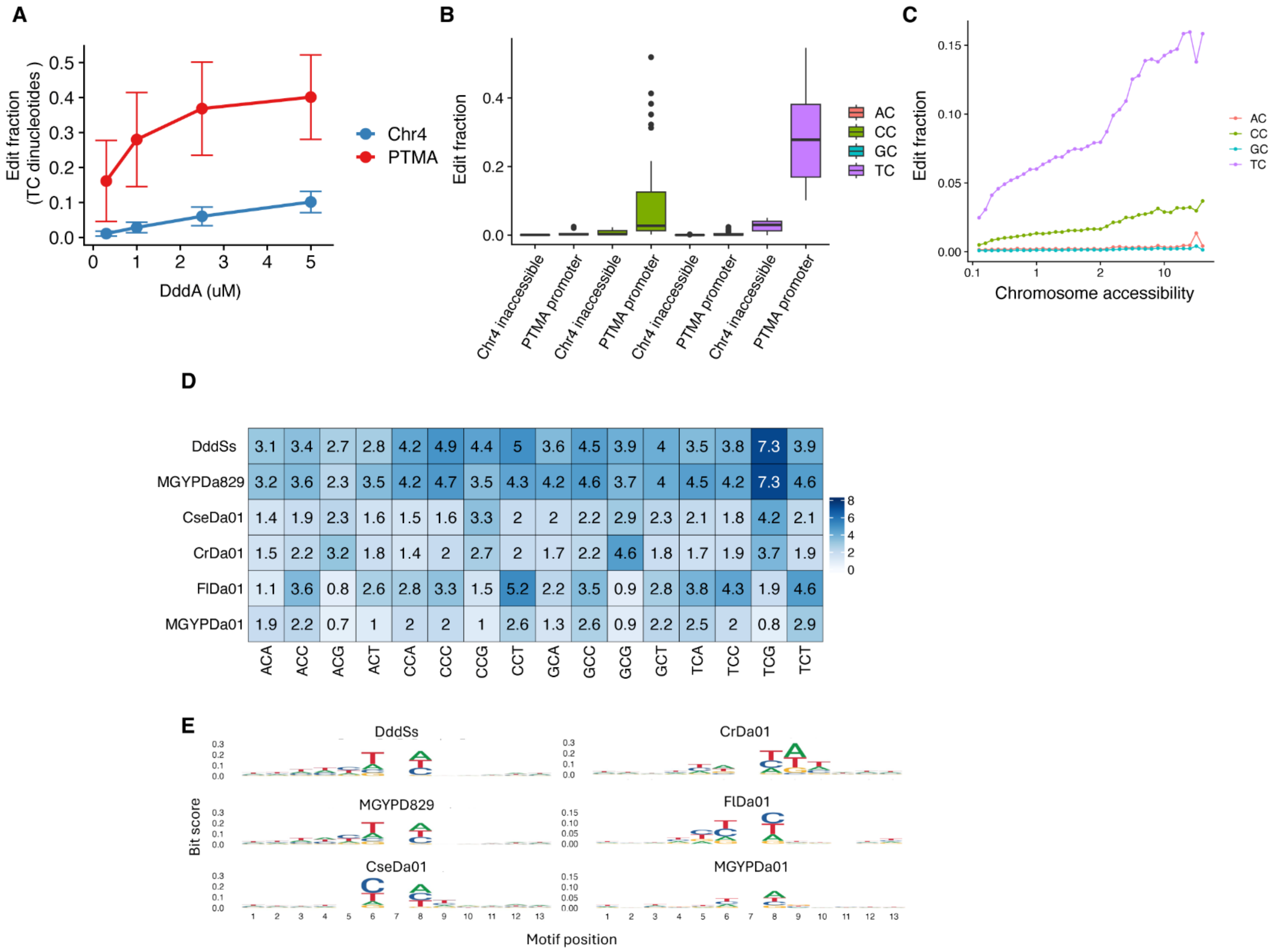


| | ACA | ACC | ACG | ACT | CCA | CCC | CCG | CCT | GCA | GCC | GCG | GCT | TCA | TCC | TCG | TCT |
|---|---|---|---|---|---|---|---|---|---|---|---|---|---|---|---|---|
| DddSs | 3.1 | 3.4 | 2.7 | 2.8 | 4.2 | 4.9 | 4.4 | 5 | 3.6 | 4.5 | 3.9 | 4 | 3.5 | 3.8 | 7.3 | 3.9 |
| MGYPDa829 | 3.2 | 3.6 | 2.3 | 3.5 | 4.2 | 4.7 | 3.5 | 4.3 | 4.2 | 4.6 | 3.7 | 4 | 4.5 | 4.2 | 7.3 | 4.6 |
| CseDa01 | 1.4 | 1.9 | 2.3 | 1.6 | 1.5 | 1.6 | 3.3 | 2 | 2 | 2.2 | 2.9 | 2.3 | 2.1 | 1.8 | 4.2 | 2.1 |
| CrDa01 | 1.5 | 2.2 | 3.2 | 1.8 | 1.4 | 2 | 2.7 | 2 | 1.7 | 2.2 | 4.6 | 1.8 | 1.7 | 1.9 | 3.7 | 1.9 |
| FlDa01 | 1.1 | 3.6 | 0.8 | 2.6 | 2.8 | 3.3 | 1.5 | 5.2 | 2.2 | 3.5 | 0.9 | 2.8 | 3.8 | 4.3 | 1.9 | 4.6 |
| MGYPDa01 | 1.9 | 2.2 | 0.7 | 1 | 2 | 2 | 1 | 2.6 | 1.3 | 2.6 | 0.9 | 2.2 | 2.5 | 2 | 0.8 | 2.9 |

**Supplementary Figure 2: Analysis of editing efficiency and motif preference of Ddd enzymes**

(A) Evaluation of deamination at TC motifs in the PTMA promoter and an inaccessible region on chromosome 4 in DddA-treated HCT116 cells. Cells were treated with four DddA doses. (B) Evaluation of deamination in the PTMA promoter and an inaccessible region on chromosome 4 in HCT116 cells treated with 1 uM DddA. Plots are divided by the nucleotide preceding the target cytosine. (C) Dinucleotide-resolved edit fractions in DddA-treated HCT116 cells binned by ATAC-seq coverage. Data derive from ACCESS-WGS data. (D) Fold enrichment of cytosine editing in accessible vs. inaccessible chromatin for 6 Ddd enzymes, separated by trinucleotide centered on target cytosine. (E) Motif logos showing the relative importance of each nucleotide surrounding the target cytosine in influencing editing efficiency for 6 Ddd enzymes. Data derive from HCT116 ACCESS-WGS data.

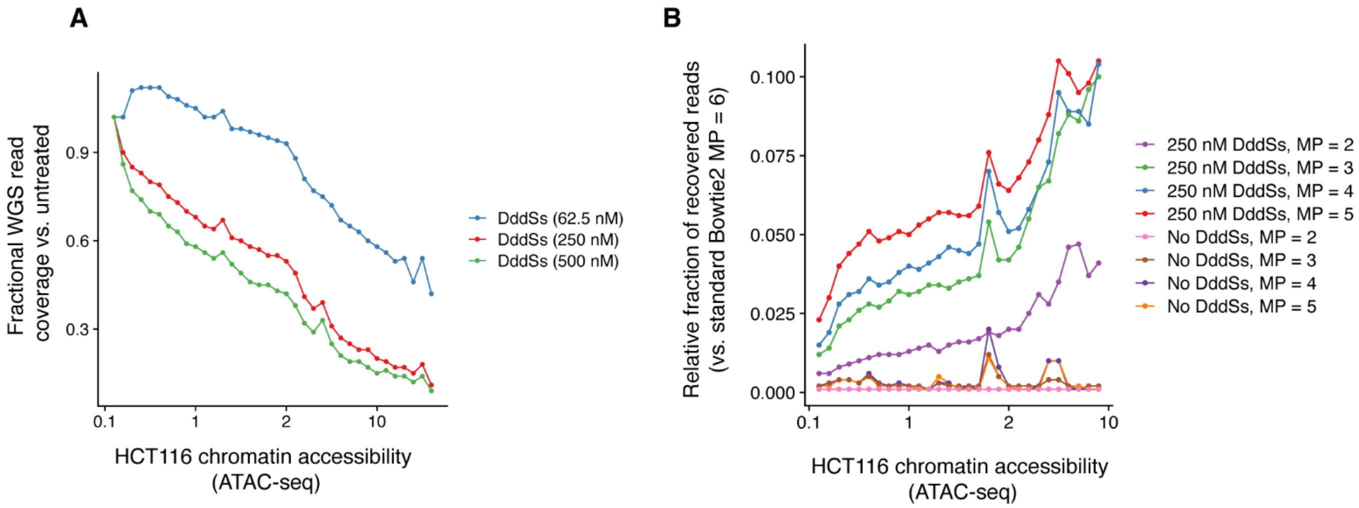


**Supplementary Figure 3: Iterative Bowtie2 alignment recovers highly edited ACCESS reads**

(A) Evaluation of relative read coverage in HCT116 ACCESS-WGS samples as compared to untreated WGS using default Bowtie2 alignment settings. DddSs treatment concentration is shown, and bins are from HCT116 ATAC-seq. (B) Evaluation of read recovery from HCT116 ACCESS-WGS samples using iterative Bowtie2 alignment. DddSs-treated and untreated WGS samples are shown at iteratively decreased mismatch penalty (MP) settings, and bins are from HCT116 ATAC-seq.

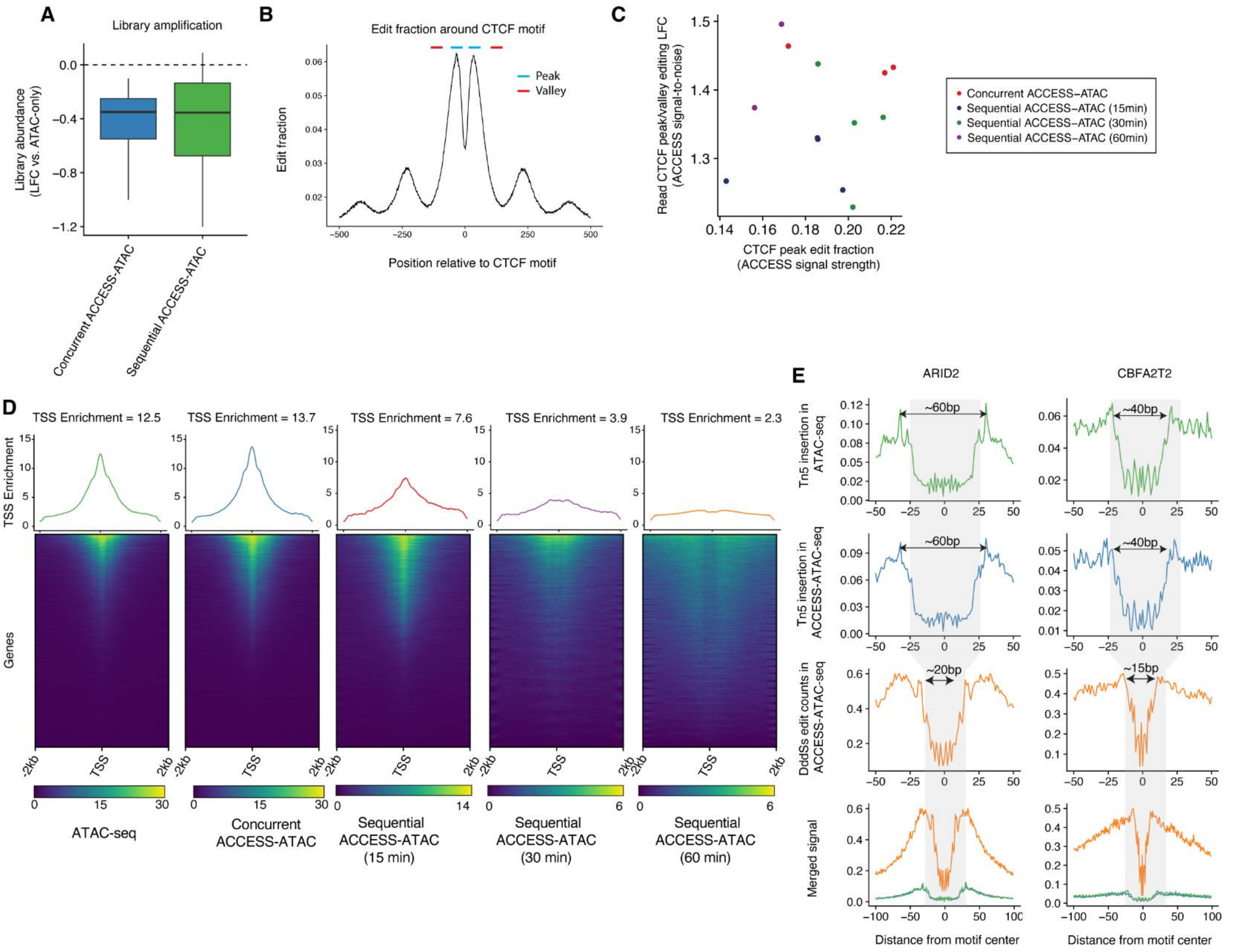


**Supplementary 4: ACCESS-ATAC combines high-resolution DddSs editing with accessible chromatin enrichment**

(A) Comparison of Ct from qPCR after Tn5 insertion step in ACCESS-ATAC vs. ATAC-seq. Concurrent and sequential ACCESS-ATAC show minimal decrease in library abundance. (B) DddSs edit fraction surrounding K562 CTCF ChIP-seq bound motifs. Peak (blue) and valley (red) regions used to calculate ACCESS signal-to-noise are represented. (C) Analysis of ACCESS edit fraction and signal-to-noise from concurrent and sequential ACCESS-ATAC protocols. (D) Enrichment of transcription start sites (TSS) in K562 ATAC-seq and concurrent and sequential ACCESS-ATAC. Each row is a TSS, rows are ordered by the magnitude of read enrichment, and a summary plot above shows read enrichment in the 2 kb surrounding all TSS. (E) Metaplots showing K562 ATAC-seq Tn5 insertion signal (green), and ACCESS-ATAC Tn5 insertion (blue) and DddSs editing (orange) signal surrounding K562 ChIP-seq binding sites for ARID2, CBFA2T2 and CUX1. Bottom plot shows merged non-normalized signal tracks to highlight the denser DddSs editing signal. Arrows denote footprint width.

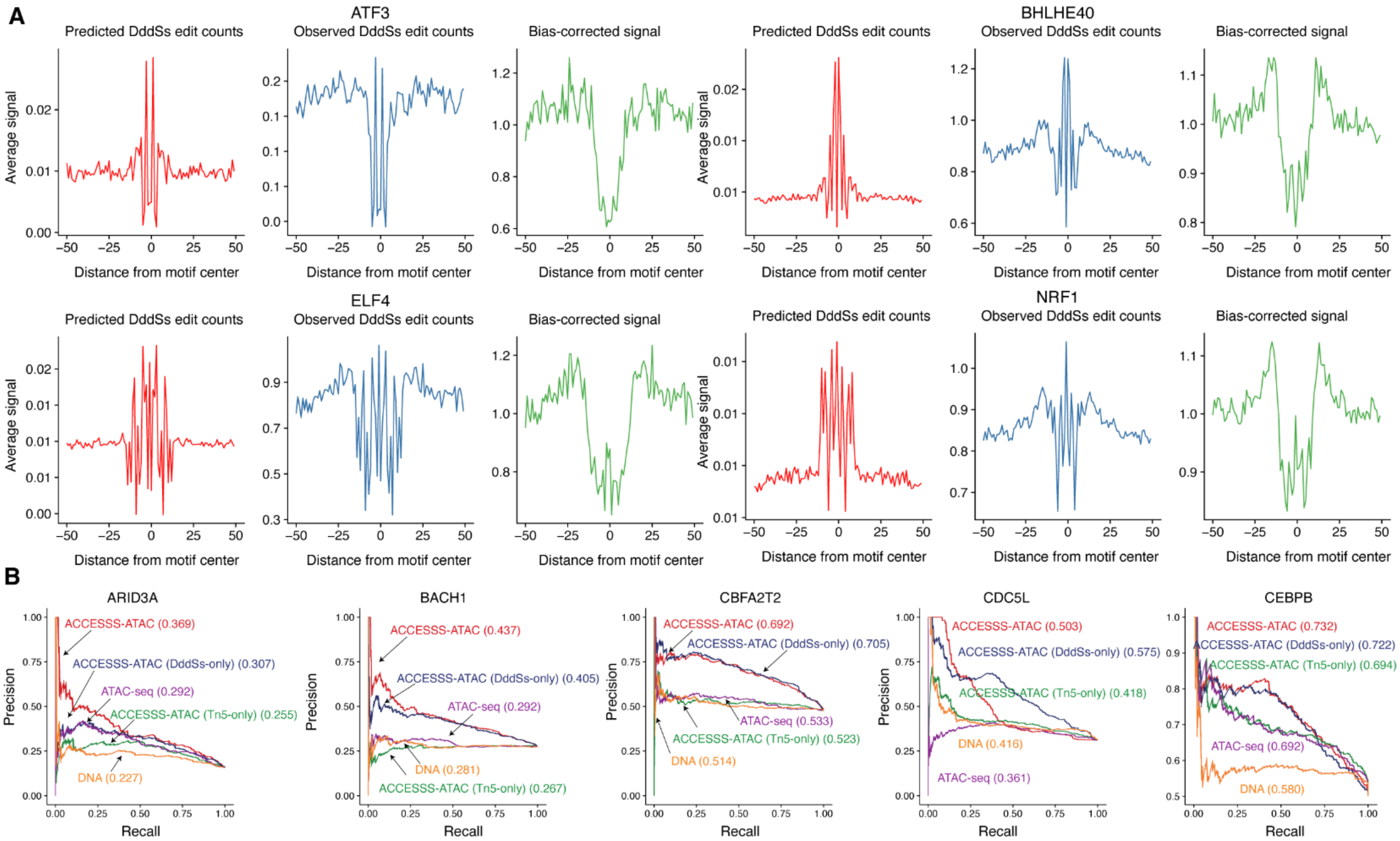


**Supplementary Figure 5: Utility of ACCESS-ATAC in predicting transcription factor binding sites**

(A) Single-nucleotide resolution signal for predicted (red), observed (blue) and bias-corrected (green) DddSs edit counts around K562 ATF3, BHLHE40, ELF4, and NRF1 ChIP-seq binding sites from K562 ACCESS-ATAC data. (B) Precision-recall curves of TFBS prediction performance for K562 ARID3A, BACH1, CBFA2T2, CDC5L, and CEBPB ChIP-seq binding sites using the indicated tracks. auPRC for each track is reported in parentheses.

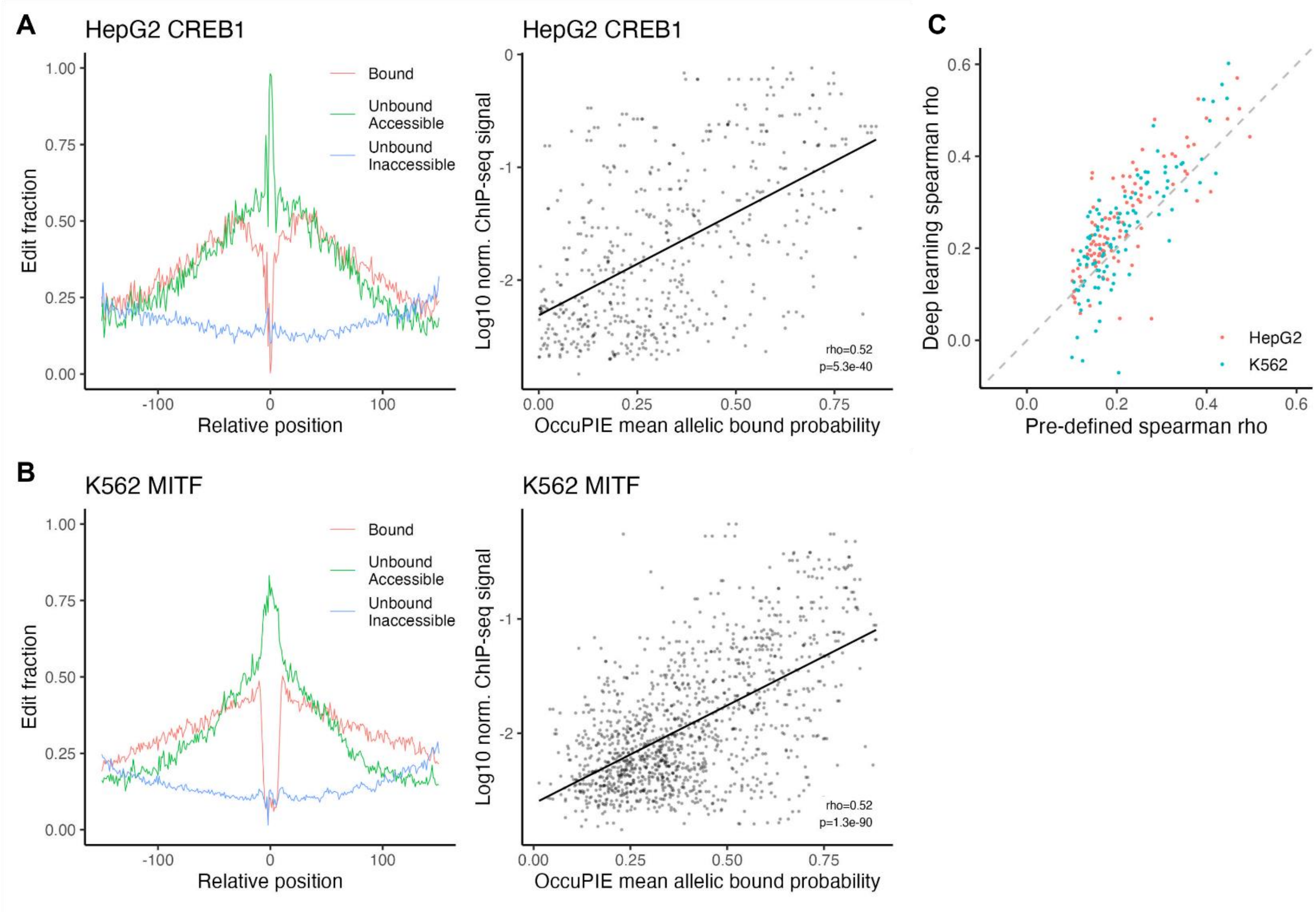


**Supplementary Figure 6: Evaluating OccuPIE, a model to impute allelic occupancy**

(A-B) Left: Edit fractions for reads surrounding K562 MITF (A) and HepG2 CREB1 (B) ChIP-seq-bound motifs. Reads are separated into bound (red), unbound, accessible (green), and unbound, inaccessible (blue) states according to maximum OccuPIE-assigned probability. Right: Evaluation of mean OccuPIE-imputed bound probability (x-axis) and ChIP-seq read abundance (y-axis) at K562 MITF (A) and HepG2 CREB1 (B) ChIP-seq bound loci. (C) Comparison of the correlation of predefined bound allele labels vs. OccuPIE-imputed labels with ChIP-seq read abundance. Spearman rho for this correlation is shown for 98 HepG2 and 106 K562 TFs.

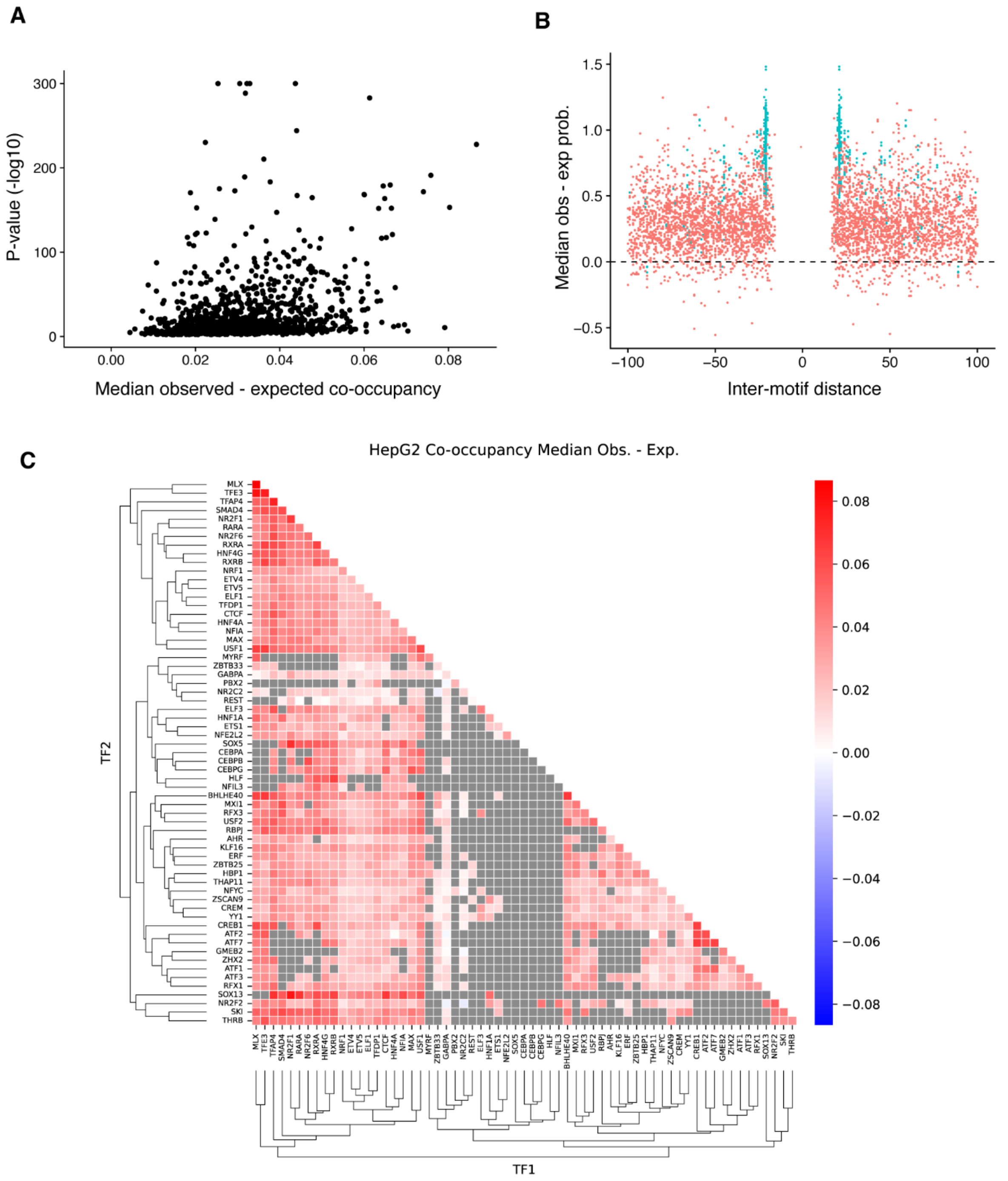


**Supplementary Figure 7: Evaluation of allelic co-occupancy of 64 HepG2 and K562 transcription factors**

(A) Evaluation of magnitude (x-axis, median observed – expected co-occupancy) and statistical significance (y-axis, $-\log_{10}(p)$) of allelic co-occupancy for 2,560 TF pairs in HepG2. (B) Heat map showing median allelic co-occupancy enrichment for all 2,560 TF pairs in HepG2. (C) Spatially resolved median allelic co-occupancy enrichment analysis of all loci within 100 nt of a K562 STAT5A motif. Heterotypic pairs (STAT5A with 63 TFs) are shown in blue, and homotypic STAT5A-STAT5A pairs in red. The high abundance of STAT5A-STAT5A pairs at 20-25-nt distance with elevated co-occupancy is notable.

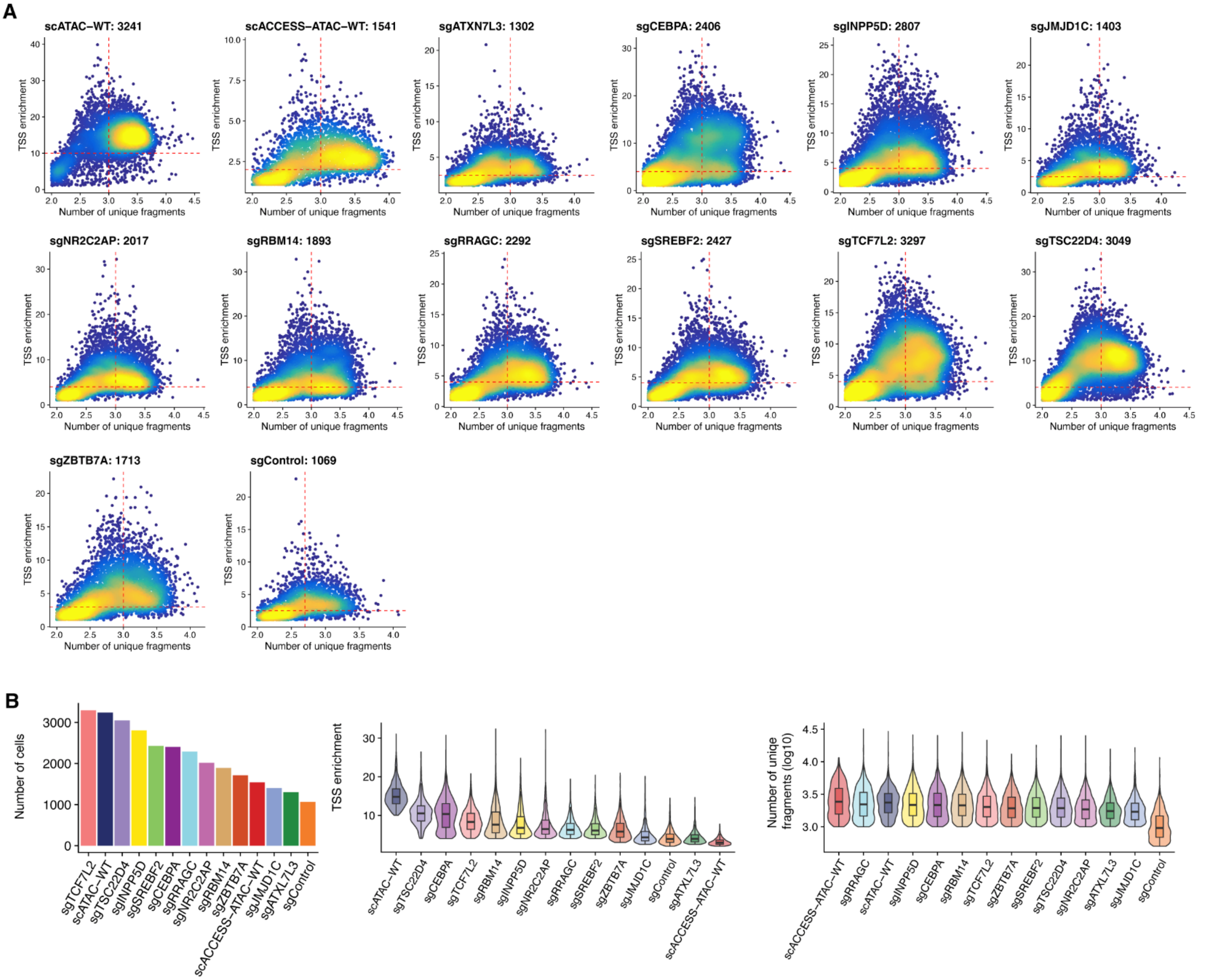


**Supplementary Figure 8: Single-cell ACCESS-ATAC data analysis**

(A) Quality control for scACCESS-ATAC data. The x-axis represents the number of unique fragments per cell, and the y-axis represents the TSS enrichment. Colors refer to the density of cells. The number of valid cells for each sample is shown at the top. (B) Left: bar plot showing the number of cells across all samples. Middle: TSS enrichment across all samples. Right: number of unique fragments across all samples.

**Supplementary Tables**

Supplementary Table 1: Motif-centered transcription factor binding prediction results

Supplementary Table 2: Motif-free footprint detection results

Supplementary Table 3: Results of allelic occupancy imputation using OccuPIE

Supplementary Table 4: Pairwise transcription factor co-occupancy enrichment

Supplementary Table 5: Transcription factor co-occupancy periodicity analysis

Supplementary Table 6: CRISPR-Cas9 gRNA oligos used in single-cell ACCESS-ATAC
